# Spatiotemporal dynamics and substates underlie emotional signalling in facial movements

**DOI:** 10.1101/2024.09.02.610622

**Authors:** Hélio Clemente José Cuve, Sophie Sowden-Carvalho, Jennifer Cook

## Abstract

From overt emotional displays to a subtle eyebrow raise during speech, facial expressions are key cues for social interaction. How these inherently dynamic facial signals encode emotion across non-verbal expression and speech remains only partially understood. In Study 1 we recorded participants’ facial movements signalling happy, sad and angry emotions in *Expression-only* and *Emotive-speech* conditions. We employed a data-driven pipeline integrating facial motion quantification, spatiotemporal classification and clustering to investigate the structure and function of facial dynamics in signalling emotion. Results reveal that a few spatiotemporal patterns reliably differentiated emotion in non-verbal expressions and emotive speech facial signals. Furthermore, we identified transient substates – or dynamic phases – that are diagnostic of emotion intent and conditions. A perceptual validation with naïve observers (Study 2) showed that the low-dimensional spatiotemporal structure captures meaningful cues that closely predict human emotion categorisations. We discuss theoretical implications of a low-dimensional spatiotemporal structure for optimal transmission and perception of dynamic facial emotion signals and face-to-face interaction. This work also provides a framework for modelling dynamic social cues and insights for the design of expressive emotive capabilities in social agents.

## Introduction

The human face is an incredibly versatile signalling tool for social interaction ^1^ capable of producing a rich array of non-verbal signals through the coordination of basic motor facial action units (AUs) ^2^. For example, frowning (AU4) combined with pursed lips (AU24) can convey a prototypical facial expression of anger, while raised cheeks (AU6) and pulling back the lip corners (AU12) can signal a smile. In addition to ostensive non-verbal emotional cues, in real-life social interaction, facial expressions simultaneously modulate verbal communication. A slight tightening of the eyelids (AU7) for example, can make a “neutral” verbal utterance appear slightly angrier. However, while much of the contemporary theoretical debate continues to focus on how specific AUs signal emotion (see ^3–5^ for a review), the mechanisms that afford such flexible expressive capabilities to facial signals remain only partly understood.

One overlooked aspect in previous research is the dynamic features of facial expression signalling, with a large focus instead on the static aspects of facial behaviour or approaches that ignore temporal dynamics ^6^. Moreover, although most face-to-face interactions naturally involve speech, the role of speech-related facial dynamics in emotion signalling has been largely neglected. Yet these dynamic cues in conversational or speech-related facial expressions can enhance the simultaneously transmission of visual signals for emotion and verbal communication ^6–9^. Nonetheless, there is growing research providing important insights on the role of facial expression dynamics to emotion signalling. Psychophysics inspired approaches have combined generative synthesis of facial movements ^10^ and reverse correlation to parametrically map the spatiotemporal features of AUs to perceptual representations of emotion categories^11^, valence-arousal dimensions ^12^ and even social traits^13^. Similarly, a recent production and perception study recorded participants’ facial movement signals of anger, happiness, and sadness, during expression only production and during spoken sentences conveying the same emotions ^14^. Kinematic analysis revealed faster movements for high-arousal emotions (happiness, anger) compared to slower movements for low-arousal emotions (sadness). Thus, suggesting that dynamic features of facial signals carry fundamental diagnostic spatiotemporal information for differentiating emotional expressions at both the production and perception stages.

Nevertheless, an account of the spatiotemporal structure of facial signals – how AUs dynamically combine and unfold to signal emotions and simultaneously convey speech-related information – remains lacking (see ^1,15^). One significant challenge to studying the role of spatiotemporal structure in facial expression signalling is the lack of an integrated theoretical and methodological framework. Such a framework should ideally integrate the dynamic nature of facial expression signals with their socio-communicative functions and provide an appropriate modelling strategy – see ^1,10^. In this work, we build on the observation that facial expressions are controlled by an underlying motor system ^16,17^. Specifically, we explore the application of theoretical and analytical principles drawn from the motor control literature to understand the spatiotemporal structure and function of dynamics in facial expressions for signalling emotion.

A motor control perspective provides two useful theoretical insights that may be applicable to studying the spatiotemporal structure and function of facial expression dynamics. First, a longstanding motor control idea suggests that body movements are generated by a low-dimensional mechanism that underpins their effective neural control and physical execution ^18–20^. Such low-dimensionality of motor control and execution might reflect the brain’s solution to the problem of controlling numerous muscles and components of the motor system whilst minimising computational demands - often referred to as the “degrees of freedom” problem ^21–23^. Such low dimensionality has been shown to describe motion patterns during walking ^24^, whole body pointing ^25^, eye movements ^26^ and articulatory speech movements ^27^. Crucially, these movements also convey emotion-related information ^28–30^ providing theoretical basis and experimental evidence for the potential of shared control principles for social and non-social aspects of movement. Similarly, dynamics underlying facial expressions, and their ability to multiplex information (e.g., visually signal emotion and speech), may be structured around low-dimensional spatiotemporal patterns. Such structure could be optimal for flexible and precise spatiotemporal control of Action Units (AUs) and to signal emotion adaptively in social interactions^31^. Additionally, given that social interaction often involve continuous verbal exchanges which impose articulatory demands to facial movements, a low-dimensional spatiotemporal mechanism may also support the simultaneous layering of emotion cues in speech-related facial dynamics^32^. This perspective also suggests a methodological framework to model spatiotemporal nature of movement, which is rare in the study of facial expression production. For instance, spatiotemporal dimensions can be explored using data-driven models that learn the underlying latent patterns of dynamic facial behaviour ^33^.

A second insight from the motor-control literature stems from the empirical observation that although outwardly fluid, body movements can often be segmented into distinct ’substates’ often seen in differential modulation of kinematics (e.g., speed, displacement of motor actions). For instance, walking movements can be segmented into ’swing’ or ’stance’ phases ^24^ or eye movements into ’fixations’ and ’saccades’ ^34^, and even more fine-grained states (sub movements) have been described ^35^. As a class of motor actions, facial expressions are likely to exhibit analogous substates shaped by the biomechanics of how they are produced and controlled, encompassing ’transitions’ where AUs are initiated or inhibited, and ’stability’ where AUs are sustained ^31^. Indeed, previous descriptions of facial motion and AU dynamics have noted such temporal segmentation during expressions ^36,37^ and emotive speech ^32^. However, substates of facial expressions and emotive speech-driven dynamics have not yet been formally characterised, and whether and how they relate to emotional signalling is unknown. Similar to spatiotemporal components, substates can be explored in a data-driven fashion, using methods such as spatiotemporal clustering ^33,38,39^ which is particularly suitable given that lack of *apriori* defined profiles for facial expression dynamics.

Probing the underlying spatiotemporal structure and substates in facial expressions would enable a richer characterisation of facial dynamics that is both holistic, i.e., captures overall spatiotemporal patterns, but also focal as it can capture micro spatiotemporal patterns. We hypothesise that these underlying patterns are not merely incidental; rather, they are diagnostic of emotional content, reflecting how facial movements are optimised for social signalling. In this context, diagnostic means that the spatiotemporal features provide reliable cues for inference of the emotion being expressed from both a production and a perceptual perspective. While some prior work has proposed similar motor-centric approaches to model facial expression signalling, these have largely focused on perception e.g. ^11,40^ rather than production. Notwithstanding, a few recent studies have employed spatiotemporal dimensionality reduction approaches to identify dynamic motifs in facial expression production data (e.g. ^22,41,42^). Nonetheless, the reliance on only a single to a handful signallers or actors limits the dynamic range and variability the facial signals analysed in prior work. Hence, it’s unclear whether the observed dimensionality in these studies, genuinely reflect the underlying mechanisms for facial expression signalling or simply results from restricted variability in signal dynamics. It is also unclear how well these findings generalise to other emotional expressions and conditions (e.g., emotive speech facial dynamics). Furthermore, previous work also relied on averaging across the spatiotemporal dimensions when comparing expressions and conditions, which re-introduces the problem of overlooking how features of facial dynamics relate to emotion signalling (see Methods). Consequently, the functional role of facial dynamics (both low-dimensional spatiotemporal patterns and substates) remains only partially explored.

The current study therefore aimed to (1) describe the spatiotemporal structure underlying facial expression production; (2) explore and characterise putative facial expression substates, and 3) evaluate the diagnostic value of spatiotemporal structure and substates for emotion signalling. To capture wider facial dynamics relevant for social interaction, in study 1 we recorded 43 participants facial movements while they signalled happy, sad and angry emotions in a) *Expression only* and b) *Emotive speech* conditions across two separate sessions, building on validated facial expression production paradigms ^14,32,43^. We used automated Facial Action Coding System (FACS) analysis to describe the spatiotemporal patterns of the facial AUs (e.g., mouth widening, eyebrow raise, etc). We then employed a novel data-driven pipeline combining dimensionality reduction, dynamic feature extraction, and spatiotemporal classification and clustering; to identify diagnostic latent spatiotemporal patterns (groups of co-activated AUs and salient temporal patterns) underlying optimal emotion signalling through facial dynamics - see Methods.

To precis our results, we find that *Expression only* and *Emotive speech* facial dynamics can be distilled into a of small set of “fundamental” spatiotemporal components. The three emotions (happy, angry, sad) and two conditions (*Expression only* and *Emotive speech*) studied are differentiated by synergistic mixing of components and subtle local substate differences related to expression transitions. In Study 2, an independent perceptual validation with naïve observers showed that the low-dimensional spatiotemporal structure is strongly predictive of human emotion categorisations.

Our key contributions are two-fold. First, we provide new insights into the complex and flexible signalling of emotion through facial movements, suggesting that it is achieved through variations in a small number of functional and perceptually relevant spatiotemporal patterns and substates. Second, we provide a versatile data-driven pipeline to model the structure and function of facial dynamics in emotion signalling, which can be extended to other dynamic social signalling modalities.

## Results

### Facial expression production, perceptual validation check

A perceptual validation of the facial expression production task recordings was conducted with 45 naïve participants in a separate study. Results show that the perceptual ratings of emotion are reliably aligned with the predicted facial expression signals from the facial production task. Variability in perceptual ratings indicate that the facial expression recordings contain sufficient diversity, with no evidence of ceiling or floor effects. Full details and figures for this validation are provided in the SI - “Perceptual Validation Study” and Fig. S1).

### Spatiotemporal components for facial expression production

We report results for *Expression only* and *Emotive speech* expressions separately for clarity, however we note that these patterns were consistent when analysed together (see SI - Comparison of joint vs separate spatiotemporal analysis of *Expression only* and *Emotive speech*, and Fig. S8-9).

#### Spatiotemporal patterns during facial *Expression only* condition

We first looked at the spatiotemporal components identified via a Non-Negative Matrix Factorization (NMF) analysis, which represents the underlying (reduced) diagnostic spatiotemporal patterns (e.g., groups of AUs with common activation timecourses and salient temporal patterns). The optimal number of spatiotemporal components (*k*) optimising for various fit indices (see Methods) – was consistently three (see Fig 1 – top panel) (see also *Expression only* condition - NMF validation analysis in Supplementary Information(SI) – Fig S2-4).

**Figure 1.**
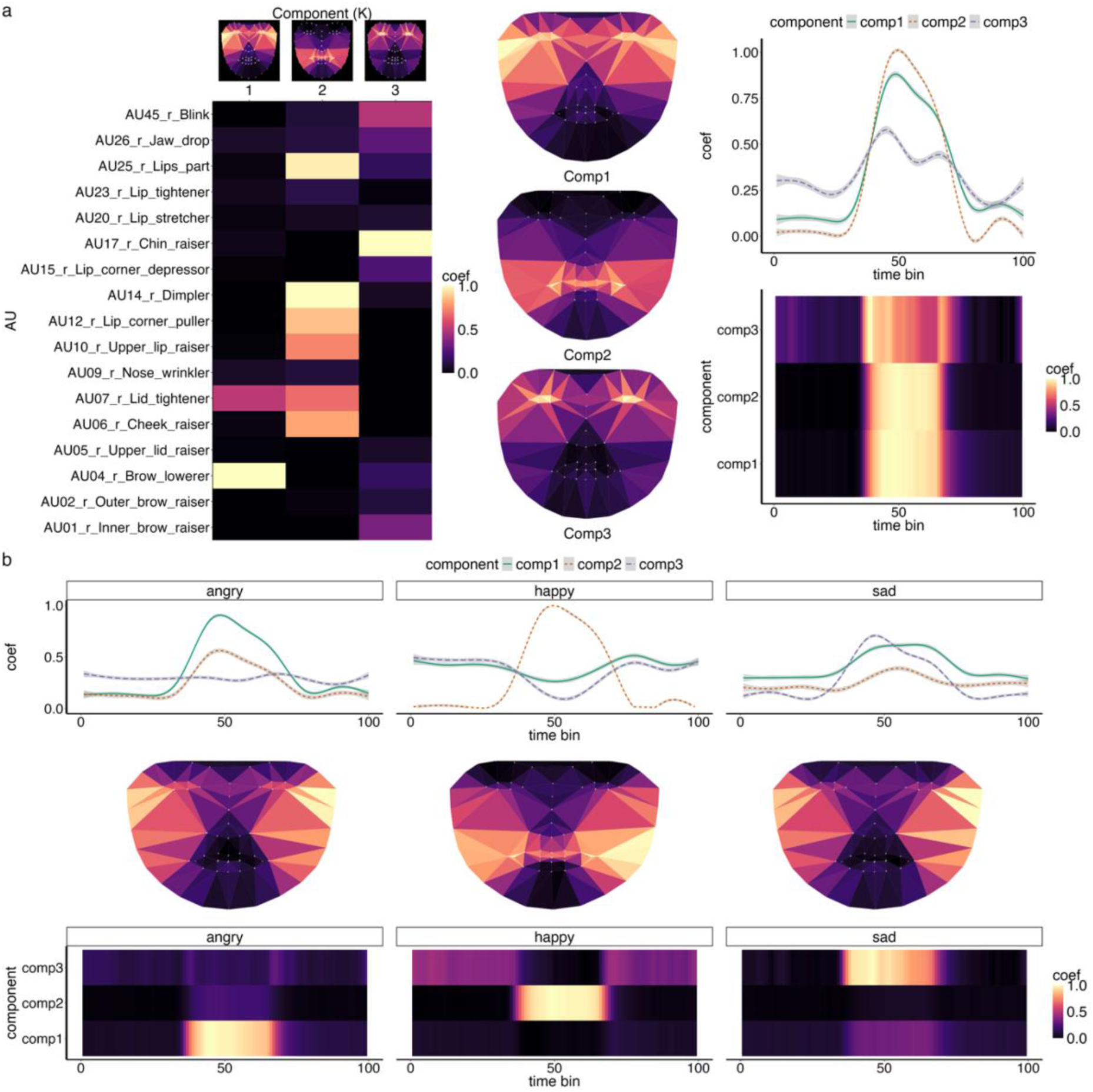
Spatiotemporal components for facial dynamics in *Expressions only* production. The top panel (a) shows that expression production can be summarised by three components, capturing key spatiotemporal patterns for upper face AUs (component 1), lower face AUs (component 2) and a combination of both lower and upper face AUS (component 3). Top - middle and right columns show approximate spatial distribution of components in an average face model and how it changes over time, respectively. The bottom panel (b) highlight spatiotemporal profiles (averaged) for angry, happy and sad expressions. Coef = Coefficient, brighter colours indicate stronger activation. Note that face maps are drawn for visualisation purposes only to provide a rough visual representation of the AU-component profiles. Faces maps are derived from an approximate pls regression predicting aligned and normalised facial landmark changes from AUs based on facial production data (see methods).

Evaluation of the spatiotemporal patterns revealed that component 1 (C1) had stronger activation of the upper face-related AUs (e.g., AU4 brow lowerer, AU7 lid tightener) compared to other AUs. Component 2 (C2) had strong activation of both lower and upper face-related AUs (e.g., AU25 lips part, AU14 dimpler). Component 3 (C3) was mostly dominated by lower face actions (e.g., AU17 chin raiser) with some small activation of upper face AUs (e.g. AU45 blink). For clarity of presentation, we name these components, upper face AUs (C1), lower face AUs (C2) and lower-to-upper face AUs (C3). The averaged temporal profile of these components across videos is shown in Fig 1 B. C1 (upper face AUs) and C2 (lower face AUS) had a common inverted-U shape with a clear singular peak, whereas C3 (lower to upper face AUs) had a less pronounced but more dynamic pattern (e.g. bimodal peaks).

Next, we formally assessed the diagnostic value of the spatiotemporal components, by analysing how features capturing dynamics differed between expressions and validated it in a held-out test set (see Methods - *Multivariate Timeseries Classification*). When assessing spatiotemporal components’ ability to classify expressions we achieved significantly above chance accuracy (ACC (accuracy)= 0.91 [Bal.Acc. (balanced accuracy) = .93], Cohens Kappa (classification reliability) = 0.87; AUC(Area Under the Curve) mean = 0.98, p < .001; note chance is .33 for a 3-way classification). These results indicate strong agreement between prediction from spatiotemporal model and ground-truth (from facial expression production task) – see Fig 2 below.

**Figure 2.**
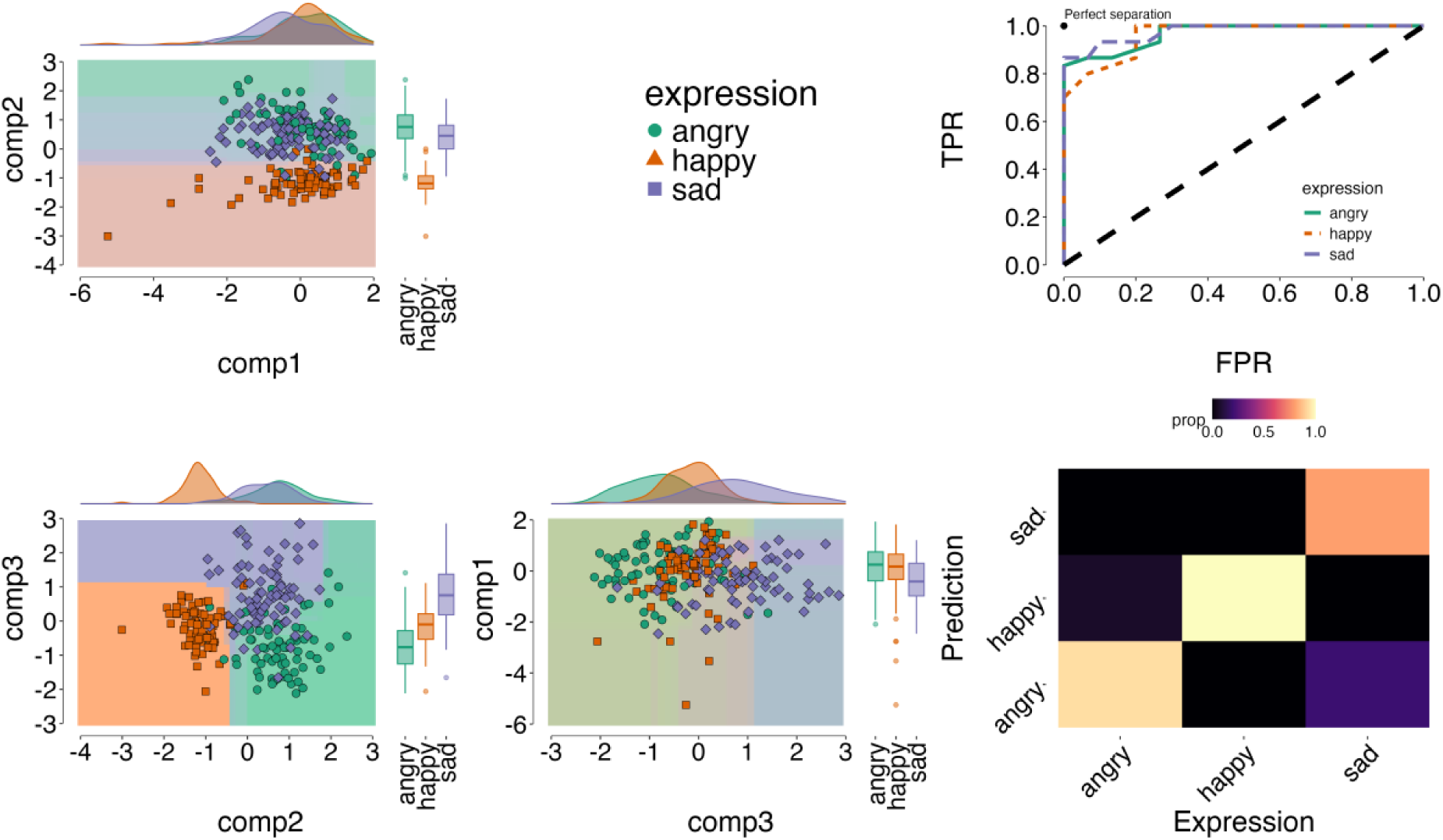
Spatiotemporal classification of facial expressions only production. *Notes:* The left panels show decision boundaries (filled backgrounds) across spatiotemporal components (comp1: upper AUs, comp2: lower face AUs; comp3: lower to upper face AUs) estimated on a training set to predict emotion classes (angry, happy, sad). Coloured dots represent individual ground-truth expressions from both training and held-out test data. Dots within a matching background indicate correct classifications, while mismatches indicate misclassifications. The results demonstrate robust classification, with at least a pair of component combinations separating a target expression from the rest, e.g. comp1 vs 2 and comp-2 vs 3 robustly distinguishes happy expressions. The top right panel presents Receiver Observer Characteristic (ROC) curves, showing good model performance (higher True Positive Rate [TPR] and lower False Positive Rate [FPR]) with the diagonal dashed line representing chance). The bottom right panel displays a confusion matrix. Brighter colours indicate higher accuracy along the diagonal elements, with off-diagonal showing little to no confusions (darker colours).

We also found distinct temporal profiles for each emotion. Angry expressions (Bal.Acc. = .91, AUC=.97, p < .001) were characterised by strong temporal coactivation of C1 (upper face AUS) and C2 (lower face AUS). Happy expressions (Bal.Acc. = .98, AUC = .99, p < .001) were dominated by C2 (lower face AUs); and sad (Bal.Acc. = .98, AUC = .90, p < .001) characterised by moderate changes in all three components. To further validate our model, we conducted a Fisher’s exact test on the confusion matrix (see Methods), which confirmed significant pairwise differentiation of all expressions on the basis of their spatiotemporal patterns (angry vs happy; angry vs sad; and happy vs sad: p.corrected < .001). These results reinforce the robustness of our classification model, demonstrating that spatiotemporal patterns capture reliable differences between emotional expressions. For detailed sensitivity analysis and metrics see SI Fig S2-S4 and Table S1-2.

#### Spatiotemporal patterns for *Emotive speech* facial expressions

Similar to the *Expression only* condition, we found that three components effectively represented the spatiotemporal patterns of facial AUs signalling emotion in *Emotive speech* (see Fig 3). However, there were some notable differences in the spatial and temporal patterns, which likely reflect the fact that, in contrast with the *Expression only* videos, the *Emotive speech* videos comprise a combination of facial expression and speech AUs. Specifically, the *Emotive speech* components comprised a mix of both lower and upper face AUs, in comparison to mostly lower vs upper face AUs for *Expression only* conditions

**Figure 3.**
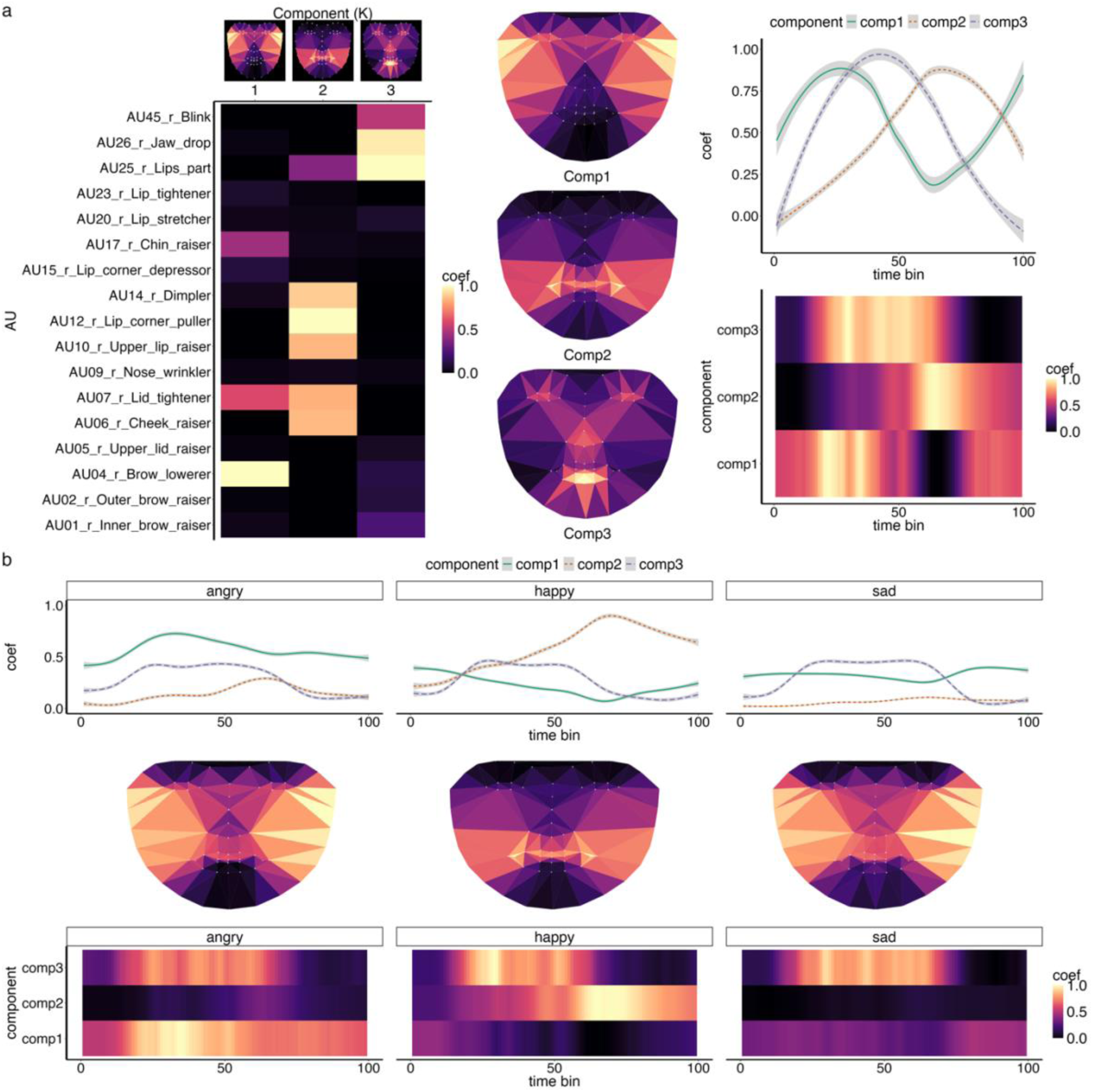
Spatiotemporal components for *Emotive speech* facial expression production. The top panel (A) shows that *Emotive speech* facial expression production can be summarised by three underlying components, capturing synergies between upper to lower face AUs (Component 1), lower-mid-upper AUs (component 2) and lower to upper face AUs (component 3). Top - middle and right columns show approximate spatial distribution of components in an average face model and how it changes over time, respectively. Note patterns are averaged across videos for visualisation. The bottom panels (B) show how components change over time, highlighting different profiles for angry, happy and sad expressions. Coef = coefficient from the NMF, brighter colours indicate stronger activation. Note while matrix heatmaps are a true representation of the model, face maps are not a one-to-one map and are drawn for visualisation purposes only. Face maps only provide a rough visual representation, derived from an approximate pls regression predicting aligned and normalised facial landmarks change from AU data based on facial production data (see methods).

Component 1 included strong activation of upper and lower face AUs (e.g., AU4 - brow lowerer; AU17 - chin raiser) peaking at both early and late time points. Component 2 (lower-mid-upper face AUs) combined strong activation of AUs in the lower (e.g., AU25 - lips part), middle (e.g., AU14 - dimpler) and upper face areas (e.g., AU7 - lid tightener). Component 3 primarily featured activation of lower face AUs, such as AU25 (lips part) and AU26 (jaw drop), plus additional moderate activation for AU45 (blink). Henceforth component 3 is referred to as lower to upper face AUs (see Fig 3). C2 (lower-mid-upper face AUS) and C3 (lower to upper face AUs) had an inverted-U shape with C3 peaking earlier than C2 (Fig 3C).

As with facial *Expression only* production, *Emotive speech* expressions were also characterised by distinct spatiotemporal patterns. That is, a classifier trained on timeseries features of the spatiotemporal components (e.g., curvature, complexity) achieved significant accuracy in classifying expressions on a held-out test set, with performance at 70% (ACC = 0.69; Bal.Acc. = .76, Kappa = 0.53, p < 0.001). These results indicate moderate to strong agreement between prediction and ground-truth emotional expression labels (see Fig 4). Class-wise, the spatiotemporal components significantly distinguished all expressions. Happy had the highest accuracy (Bal.Acc. = 0.91, AUC = 0.97, p < .001), suggesting a clear and distinct spatiotemporal profile, notable in the increased activation in C2 (lower-mid-upper face AUs) over time. For detailed validation and statistics see SI Fig S5-S7 and Table S3-4

**Figure 4.**
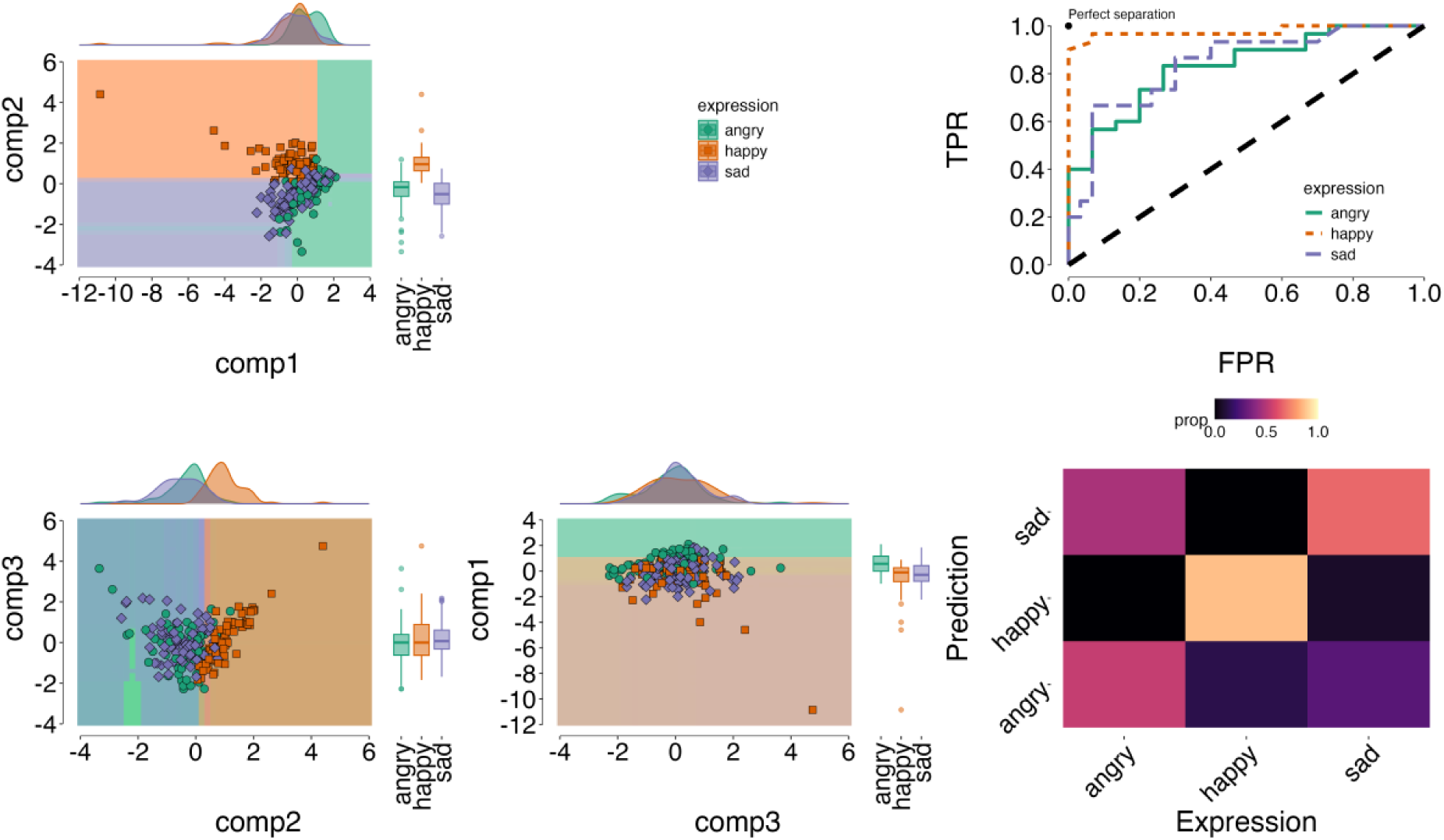
Spatiotemporal classification of *Emotive speech* facial expressions. The left panels show decision boundaries (filled backgrounds) across spatiotemporal components (comp1: upper to lower face AUs, comp2: lower to upper face AUs; comp3: upper-to-lower face AUs) estimated on a training set to predict emotion classes (angry, happy, sad). Coloured dots represent individual ground-truth expressions from both training and unseen test data. Dots within a matching background indicate correct classifications, while mismatches indicate misclassifications. The results demonstrate robust classification, with component combinations effectively separating the expressions in at least one combination. The top right panel presents ROC curves, showing good model performance (higher True Positive Rate [TPR] and lower False Positive Rate [FPR]), with the diagonal dashed line representing chance. The bottom right panel displays a confusion matrix. Brighter colours indicate higher accuracy along the diagonal elements, and confusion off-diagonal (e.g., happy shows high accuracy and low confusion, whereas angry shows higher confusion than all expressions).

Angry had the lowest classification accuracy compared to all other expressions, yet still significantly above chance (Bal.Acc. = 0.67, AUC = 0.83, p = .01), reflecting a more complex and less distinct pattern across all components. Similar to happy, sad expressions were classified reliably above chance (Bal.Acc. = 0.72, AUC = 0.84, p = 0.001). Fisher’s exact test on the confusion matrix confirmed significant pairwise differentiation between happy and sad expressions (p < .001) and between angry and happy (p < .001). However, it suggested confusions between and angry and sad (p = 0.12) (see Fig 4; SI1 - Table S3-4).

In summary, we found three latent spatiotemporal components that reliably described the dynamics of facial expressions during *Expression only* and *Emotive speech* conditions. Robust train-test classification on unseen held-out data, indicates that spatiotemporal components carry fundamental diagnostic emotional cues with good generalisability.

### Perceptual validation of the spatiotemporal components for emotion signalling

We also conducted an independent perceptual validation with 45 naïve observers who judged point-light (PLFD) animations isolating facial motion of the produced stimuli, following a validated protocol from our previous work ^14^ (See methods and SI). Results showed that the low-dimensional spatiotemporal structure is strongly predictive of human emotion categorisations in the holdout test-set. Specifically, we found (a) strong correspondence between our model’s spatiotemporal classification predictions and human observers’ categorisations (*Expression only,* X² = 33.125, p < .001, *Emotive Speech*, X² = 15.622, p = .002) and (b) high predictive accuracy of human observers’ emotion categorisation based solely on these spatiotemporal components (Accuracy consistently above 60% across stimuli and participants, and conditions; all p <. 001). For full details and statistics see SI: Perceptual validation of the spatiotemporal components for emotion signalling; and Fig 5).

**Figure 5.**
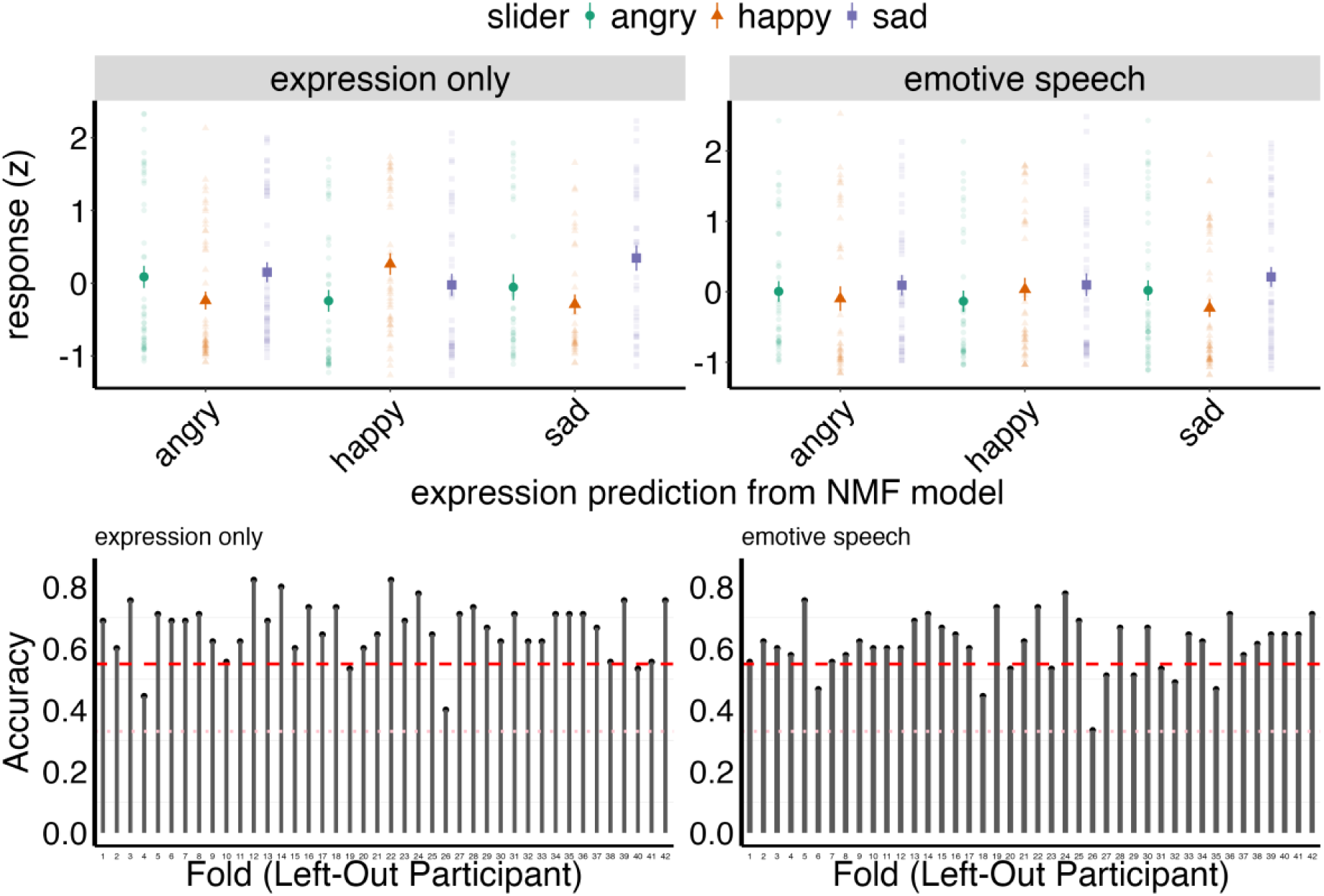
A. Mean standardised ratings for *angry* (green), *happy* (orange), and *sad* (purple) expression in the *Expression only* (left) and *Emotive speech* (right) conditions. The horizontal axis reflects the expression predicted by the spatiotemporal model. Each point indicates a single video’s average rating across participants. Data shows that the model predictions differentiate emotion ratings, despite some variability. **B.** Leave-one-participant-out cross-validation accuracy for the *Expression only* (left) and *Emotive speech* (right) conditions. Each bar shows the accuracy when a single participant’s data are held out of training. The red dashed line indicates the empirical chance level (i.e. majority class prediction), while the pink dotted line represents the theoretical chance level for a three-way classification.

### Substates reveal emotion diagnostic transitory patterns in facial signals

The second objective of this study was to identify and characterise facial expression substates. While these are likely driven by articulatory and biomechanical nature of activating AUs, we aimed to explore whether they are additionally modulated by emotional intent (i.e., they may differ between happy, angry and sad expressions and conditions). We identified three primary substates through spatiotemporal clustering that segmented periods with robust local dynamics within the underlying spatiotemporal components (see Methods: *Spatiotemporal clustering for substates analyses*). Visual representation of the kinematic profiles of each substate (Fig 6A-B) suggest the substates correspond to relaxed, transition, and sustain periods for spatiotemporal components. The overall pattern of substates is also consistent across participants, suggesting that this is not an artifact of methodology or driven by specific inputs (see Fig 6 C-D).

**Figure 6.**
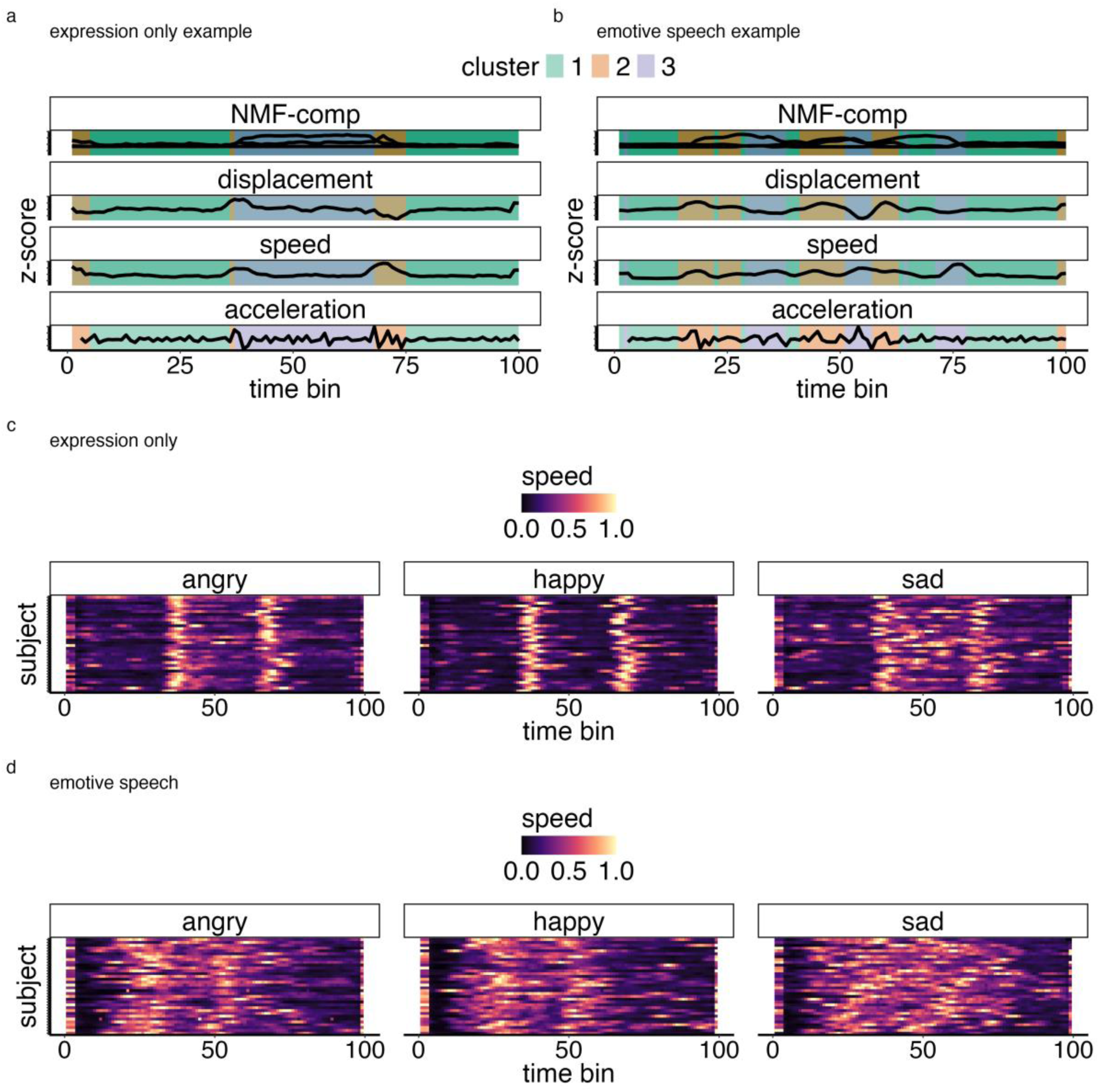
Illustration of facial expression substate profiles. **A-B.** Example of substate clusters (1 (green) = relaxed, 2 (peach) = transition; 3 (purple) = sustain). C-D show overall the speed distribution for *Expression only* and *Emotive speech* expressions for all subjects.

Because the clustering algorithm returns only cluster partitions, it does not inherently explain why these clusters emerge or how they may differentiate emotion (anger, sadness and happy) and conditions (*Expression only vs Emotive speech*). To address this, we quantified cluster (substate) patterns using two metrics, which were selected following exploratory visual inspection of the substate profiles (see Fig. 6): (1) speed as the input to the clustering analysis, provides a kinematic characterisation of each substate; (2) entropy, which provides a view of overall dynamics of substates (e.g., how they change over time) (see Fig 6 and Methods). Similar measures have also been shown to capture diagnostic information in facial expression movement and AU actuation ^11,14^. As this analysis is intended to characterise substates clusters rather than classification per se, we used a linear mixed-effects model - LMM (see Methods) to statistically explore differences in these metrics between emotion and conditions. Our overarching hypothesis, outlined in the Introduction, was that substates would differentiate emotional signals in line with the idea that local facial dynamics optimise the motor encoding of emotion cues. However, no specific directional effects were predicted a priori, as the substates were derived in a data-driven manner, and as such the results should be interpreted as exploratory, aimed at characterising the substate–emotion patterns.

#### Transition substate speeds differentiate emotional expressions

An LMM used to predict speed from emotion (happy, angry, sad), condition (*Expression only, Emotive speech*), and substate (relaxed, transition, sustain) showed strong total explanatory power (conditional R2 = 0.6 (66%)) with the fixed effects alone (marginal R2) accounting for 0.63 (63%) of the variance. All main effects were significant. First, there was a small significant effect of emotion (F(2,1990) = 12.89, p < .001). Pairwise tests showed that angry expressions were significantly slower than happy expressions (angry < happy; beta = -0.028, SE = 0.008, t(1207) = -3.398, p.corrected = 0.002). Happy expressions were faster than sad expressions (happy > sad; beta = 0.041, SE = 0.008, t(1207) = 4.92, p.corrected < 0.001). There were no significant differences in speed between angry and sad expressions (beta = 0.012, SE = 0.008, t(1207) = 1.53, p.corrected = 0.38). There was a significant main effect of condition (F(1, 1190) = 369.41, p < .001) with *Expression only* condition moving faster than spoken (beta = 0.13, SE = 0.007, t(1207) = 19.08, p < .001). Finally, there was a main effect of substate (F(2, 1190) = 687.90, p < .001) where sustain had higher speed than relaxed (beta = -0.16, SE = 0.008, t(1207) = -19.52, p.corrected < .001); transition had higher speed than relaxed (beta = -0.307, SE = 0.008, t(1207) = -36.80, p.corrected < .001) and transition had higher speed than sustain (beta = -0.144, SE = 0.008, t(1207) = -17.28, p.corrected < .001). In summary, facial movement speed varied by emotion, condition, and substate, with happy expressions, the Expression only condition and transition phases being the most dynamic overall.

All two-way interactions were significant. First, there was a significant interaction between emotion and substate (F(4, 1190) = 15.24, p < .001). Pairwise tests showed that differences in speed between happy, angry and sad emotional expressions were statistically significant for the transition substate only (happy>anger, beta = -0.077 SE = 0.014, t(1207) = -5.36, p.corrected < .001; sad< angry, beta = 0.053, SE = 0.014, t(1207) = 3.64, p.corrected < 0.001; sad< happy, beta = 0.130, SE = 0.014, t(1207) = 9.005, p.corrected < .001). There were no speed differences between emotions for the relaxed substate (angry vs. happy, beta = -0.005, SE = 0.014, t(1207) = -0.348, p.corrected = 1.0; angry vs. sad, beta = 0.007, SE = 0.014, t(1207) = 0.498, p.corrected = 1.0; happy vs. sad, beta = 0.012, SE = 0.014, t(1207) = 0.846, p.corrected = 1.0) nor the sustain substate (angry vs. happy, beta = -0.003, SE = 0.014, t(1207) = -0.177, p.corrected = 1.0; angry vs. sad, beta = -0.022, SE = 0.014, t(1207) = -1.501, p = 0.40, p.corrected; happy vs. sad, beta = -0.01913, SE = 0.0144, t(1207) = -1.325, p.corrected = 0.5567). These results suggest that transition substate (but not relaxation or sustainment) carry diagnostic information about the emotional content of facial expressions (Fig 7).

**Figure 7:**
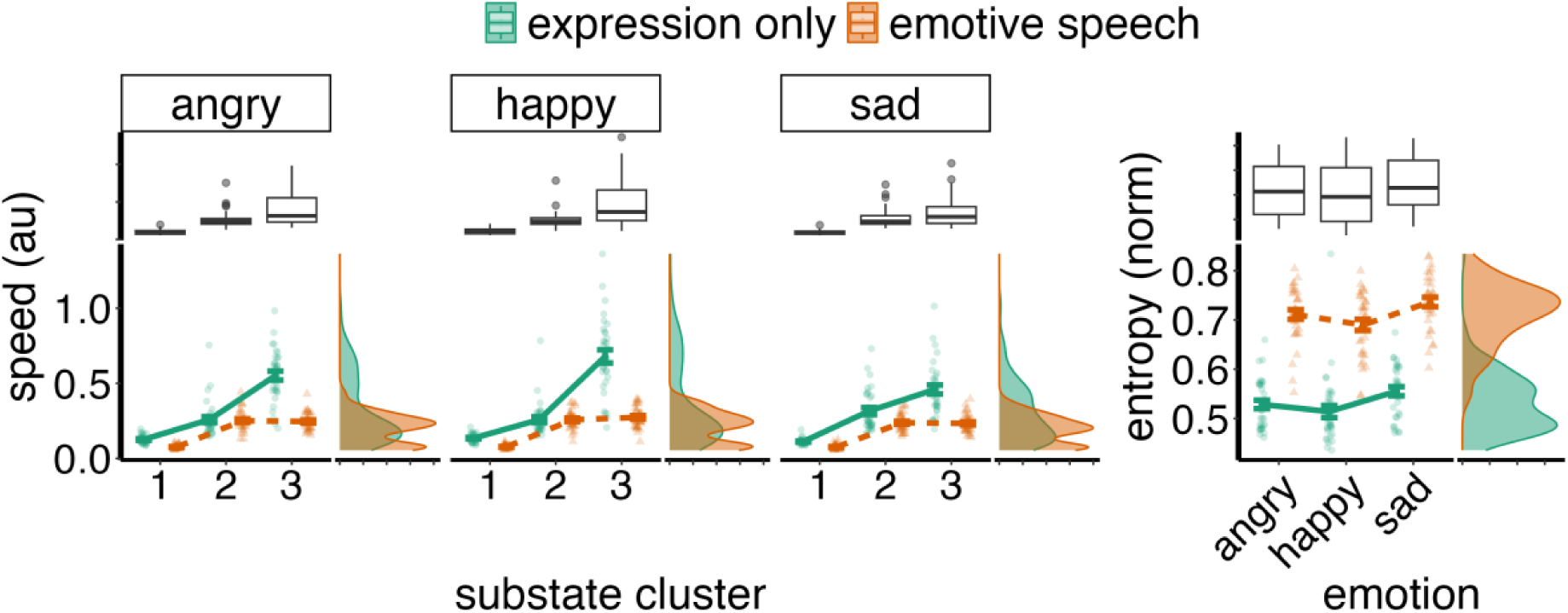
Summary of speed and entropy of substate patterns. Substate cluster: 1-relaxed, 2-sustained, 3-transition. *Expression only* condition has faster than *Emotive speech*, with transition substates being faster; Entropy mainly differentiates *Expression only* and vs*. Emotive speech* condition, with the latter exhibiting more complex patterns. Entropy also differed between happy and all other expressions, with substate patterns being more predictable for former.

Finally, there was a significant two-way interaction between condition and substate (F(2) = 183.65, p < .001). For the *Expression only* condition, there were significant differences in speed between all substates such that sustain was faster than relaxed (relaxed < sustain, beta = -0.153, SE = 0.012, t(1207) = -12.96, p.corrected < .001); transition was faster than relaxed (relaxed < transition, beta = -0.439, SE = 0.012, t(1207) = -37.23, p.corrected < .001), and faster than sustain (beta = -0.286, SE = 0.0118, t(1207) = -24.271, p.corrected < .001). These patterns were overall similar for the *Emotive speech* condition where relaxed periods were significantly slower than sustain (beta = -0.173, SE = 0.0118, t(1207) = -14.647, p.corrected < .001) and transition periods (beta = -0.175, SE = 0.012, t(1207) = -14.82, p < .001). However, there was no statistically significant difference between sustain and transition speeds (beta = -0.002, SE = 0.012, t(1207) = -0.169, p.corrected = 1.00). In summary, substate differences were more apparent in the *Expression only* condition than in *Emotive speech*, reflecting stronger optimisation of facial dynamics for expressive signalling when speech demands are absent.

In sum, substate clusters are associated with differences in speed suggesting periods of transition, relaxation and sustainment of expressions. Interestingly, these micro-dynamic patterns differ between expressions and between *Expression only* and *Emotive speech* signals, suggesting that facial production phases may enhance the signalling of the emotional intent during speech.

#### Substate dynamics differentiate emotion content

Here we used entropy as to statistically describe the dynamics of facial expression substates in terms of how temporally structured/predictable they are (^44^. see Methods). High entropy equates with high complexity in substate patterns, and low entropy with low complexity. Fig 6, for example, illustrates that substate transitions are more complex (i.e., high entropy, less predictable) in the *Emotive speech* condition where visual pattern differences between emotions are harder to discern than in the *Expression only* condition (see Methods: *Spatiotemporal clustering for substates analyses*). Note entropy was log transformed to correct for non-normal residuals.

The entropy model explained 67% of the variance (R² = 0.67). The fixed effects alone (emotion, and expression condition) accounted for 57% of the variance (marginal R² = 0.57). The main effect of emotion was statistically significant (F(2, 340) = 12.27, p < 0.001). Pairwise contrasts revealed that sad expressions had a significantly higher entropy (more unpredictable substate sequences) compared to happy expressions (estimate = -0.027, SE = 0.0055, t(344) = -4.897, p < 0.0001). There were no statistically significant differences in entropy between angry and happy expressions (estimate = 0.011, SE = 0.006, t(344) = 1.996, p = 0.14), but there was a significant difference between angry and sad expressions (estimate = - 0.016, SE = 0.0055, t(344) = -2.90, p = 0.012) – see Fig 7.

The largest effect was the main effect of condition (F(1, 34) = 429.10, p < 0.001), with the *Expression only* condition demonstrating significantly lower entropy than *Emotive speech* expressions (β = -0.208, SE = 0.017, t(35) = -12.594, p < 0.001). This suggests more complex movement dynamics in the *Emotive speech* compared to the *Expression only* condition (see heatmaps in Fig 6). There was no significant interaction between emotion and condition (F(2, 340) = 0.03, p = 0.970).

In summary, *Expression only* substates (relaxation, transitions, and sustainment of expressions) are more temporally structured than those in *Emotive speech* expressions, likely reflecting the increased dynamic complexity associated with facial movements during speech. Substates patterns also differentiate expressions, with sad expressions having a more complex dynamics than happy and angry expressions.

To summarise, in addition to the low-dimensional spatiotemporal patterns that are diagnostic of emotion and expression condition, we also identified substates reflecting periods of relaxation, transition, and sustainment of facial actions. These substates are distinguished by their unique kinematic and dynamic (complexity) profiles which provide cues for differentiating emotional expressions in *Expression only* and *Emotive speech* conditions.

## Discussion

The current study investigated how the underlying dynamics of facial movements relate to their expressive emotional signalling function. Building on a motor-theoretic framework we employed a data-driven pipeline - integrating spatiotemporal dimensionality reduction, timeseries feature extraction, classification and clustering; to identify diagnostic patterns underlying emotion signalling in both *Expression only* and *Emotive speech*. Our work contribute two key insights. First, we consistently identified a small number of spatiotemporal components that reliably described the time courses and covariance patterns of AUs. The identification of these components suggests that emotion signalling through facial expressions and speech-related facial cues can be achieved by just a few fundamental dynamic patterns; combinations of these dynamic patterns differentiate production and perception of angry, happy and sad expressions. Second, we showed that these spatiotemporal patterns exhibit a substate-like structure, which reflect transient periods of relaxation, transition, and sustainment of expressions. The profiles of these substates differed by emotion category as well as between *Expression only* and *Emotive speech* conditions, indicating they partly reflect the encoding of dynamic facial emotion cues.

The low dimensionality of the spatiotemporal patterns of facial expressions, that we have uncovered, is intriguing given the various combinations of AUs that can contribute to the same expressions ^6^ as well as the variability in how people produce expressions ^45^. Such low-dimensional spatiotemporal structure may reflect flexibility for signalling, as a variety of expressions can be rapidly produced (and perceived) by relying on just a few interchangeable dynamic patterns. Theoretically, such flexibility is likely beneficial in face-to-face interaction, where facial expressions may need to be quickly adapted according to interaction demands ^6,46^. Similarly, this flexibility might be necessary to accentuate emotional information during conditions which impose dynamic constraints on facial movements, such as speech^7,46^. These findings are consistent with the observation of low-dimensional spatiotemporal patterns facial ^22,42^ as well as other body movements ^21,23,25^ which also have the capacity to signal emotion information ^28,47^. This suggests the potential for shared control mechanisms for social and non-social aspects of movement including speech related facial movements.

An important question concerns whether our results are representative of facial expression signalling in general, or simply an outcome of the particular methodology we employed. For example, it could be argued that we identified a small number of spatiotemporal components because we asked participants to demonstrate relatively simplistic expressions which can be signalled by activating just a few action units. We argue that this is not the case because a) the *Emotive speech* condition was also specifically designed to dissuade participants from relying on overly simple and caricatured expressions and to introduce dynamic variability and b) compared to previous work ^22,42^ we provide a substantially wider range of signallers to capture wider idiosyncrasies and variability in facial dynamics (see SI- Fig S10). It could also be argued that our results are simply a consequence of dimensionality reduction, as any set of data can in theory be represented by a lower-dimensional structure. Yet, as we showed in a series of validation and sensitivity analyses, the components learned carry diagnostic information about emotions, which is robust to comparisons against appropriate randomised dynamic patterns and generalises to unseen data. Thus, we have not generated a “non-sense” low dimensional structure but rather a meaningful way to represent diagnostic dynamic cues for emotion signalling.

An intriguing question concerns why this low-dimensional spatiotemporal structure emerges. One possibility is that receivers only attend to a reduced subset of spatiotemporal patterns and correspondingly, signallers tend to only accentuate a subset of features to produce predictably diagnostic facial movements. Relevant to this argument, our perceptual validation indicates that there is overlap between model-based spatiotemporal emotion classification predictions, and human categorisation (see Fig 5). Moreover, the robust prediction of participants’ categorisations directly from these components suggests that these low-dimensional patterns are at least in part, functionally and perceptually relevant for signalling emotion. These results are also consistent with previous psychophysical approaches showing that when observers are presented with dozens of biologically plausible face movements, their perceptual representations suggest that they only rely on a reduced subset of dynamic patterns ^40^. However, there have not been systematic studies assessing the correspondence of these perceptual signals to those actually produced by observers. Furthermore, as human infants learn to produce and recognise facial expressions throughout a protracted period of development ^48^, it is possible that humans learn to optimise signals based on how these are optimally perceived and responded to, creating a reciprocal dynamic interaction between these processes. Thus, the low-dimensional organisation of facial expressions may reflect the optimisation of social signals according to perceptual demands and these perceptual demands, themselves, may be influenced by the low-dimensional organisation of facial expressions ^49^. An additional possibility is that there are savings (e.g., energy/computational, cognitive) to be had from producing and interpreting facial expressions according to only a small number of key informative spatiotemporal patterns.

The identification of substates in emotional facial dynamics also goes beyond previous work on facial expression production. On the one hand substates likely reflect the biomechanical features of articulating facial expressions, which inevitably result in periods of relaxation, transition and sustainment of the underlying motor units ^50,51^. Yet, the fact that the patterns of these substates vary with different expressive emotions and conditions even under the simultaneous production of the same verbal utterances, suggests that these local dynamics provide a medium for optimising the signalling of emotion.

Additionally, in real-life interaction, facial movements subserving verbal and nonverbal expressions are often seamlessly integrated. Our findings in the *Emotive speech* condition raise the possibility that verbal and nonverbal facial dynamics when signalling emotion can be integrated by differential modulation of spatiotemporal structure. For example, during speech, lip movement might covary with articulation and sound production while simultaneously, the tightening of the lips and lowering the eyebrows can accentuate the emotion signal. Indeed, the observation of mostly part-based profiles (e.g., lower vs upper face AUs) in *Expression only* conditions, contrasting the involvement of both mouth and non-mouth-related components in *Emotive speech* expressions supports this assertion. Similarly, the reduced substate differentiation and higher entropy observed in the substate patterns of *Emotive speech* may reflect the increasing demands placed on facial dynamics by speech articulation and emotional expression, resulting in more complex spatiotemporal patterns needed to differentiate emotion. However, given that social signalling is inherently multimodal - with tone of voice, vocal prosody, and facial expressions capable of conveying emotion both independently and in combination - a future challenge is to understand the extent to which these modalities act redundantly (conveying the same information) or synergistically (combining to produce qualitatively richer meaning) when signalling emotion ^1,12,52^.

There are several limitations and considerations that point towards valuable directions for future work. First, including more detailed descriptors of face movements (including head motion see ^42^), speech-optimised facial dynamics and the use of more naturalistic signalling paradigms, may require additional dimensions to capture the full complexity of dynamic emotion signalling. Similarly, while this study had no specific predictions about dynasticity of dynamics cues for individual emotions, we observed baseline differences in decoding accuracy. It would be informative for future studies to explore potential spatiotemporal features which may underlie such differences and potential emotional confusions at both the production and perceptual levels. Emerging computational and experimental approaches enabling controlled manipulation of spatial and temporal facial dynamics in real or near-real-time interactive settings offer promising routes for causal modelling of spatiotemporal mechanisms of production and perception of facial emotion signals ^53,54^. Second, a different number of substates might be discovered using physiological approaches, such as electromyography (EMG). Such approaches are sensitive to subtle motor potentials and therefore may reveal more substates, including preparatory or corrective movement impulses not apparent in FACS resolution, which may carry both invariant and variant cues for emotional signalling. Finally, our approach focuses on exhaustively describing spatiotemporal components and how dynamic features that describe facial motion may optimally encode emotion. However it does not isolate specific components, features (or groups of features) nor assess their individual diagnostic value of select AU activations. Our sensitivity analyses however, showed that models based on discrete AUs were generally less reliable in decoding production and perceptual data than those based on component structure. Even so, given that some AUs may be more dominant and exhibit a broader dynamic range, whereas others are more subtle or infrequently expressed, more systematic investigation of how different AUs dynamics contribute to emotional signalling is warranted.

Despite these considerations, our results and approach provide new insights into the dynamic mechanisms of face-to-face emotional signalling. This framework also offers a promising model for understanding clinical conditions such as autism and depression, particularly in relation to the role of atypical facial expression production in social interaction difficulties ^55,56^. These findings also have practical applications in fields such as social robotics and human-computer interaction. A significant challenge in these fields relates to endowing social agents (robots, digital agents) with convincing and adaptable repertoires of social behaviour, including facial signalling ^57^. Spatiotemporal components and substates underlying facial expression production could provide actionable building blocks without the need to painstakingly design each facial expression into the architecture of these systems.

## Conclusion

In conclusion, this study identifies a low-dimensional spatiotemporal structure and a set of substates that underpin emotional signalling and perception of *Expression only* and *Emotive speech* facial dynamics. It also provides a data-driven pipeline to support the development of theoretical accounts of the structure and function of expressive facial dynamics. This framework can be extended and compared to other spatiotemporal models of facial dynamics and modalities to further elucidate the dynamic and multimodal nature of emotion signalling in face-to-face interaction.

## Methods

### Participants

43 volunteers from the university community (24 female; mean age = 26), took part in the facial expression production study for course credit or monetary incentive. A separate sample of 45 participants from the general population, naïve to the production task, took part in a perceptual validation experiment (see SI Perceptual Validation Study). All participants gave informed consent, and all procedures of the study were approved by the School of Psychological Science Research Ethics Committee at both the University of Bristol and University of Birmingham (ref 21918). All procedures also adhered to the revised 2013 Declaration of Helsinki and relevant national and international guidelines on research involving human subjects. The dataset in the facial expression production study was collected as part of a larger project investigating facial expression production kinematics and perception – see ^14^. However, all the variables and analyses reported here are specific to this study and not discussed in any previous papers. Furthermore, data from perceptual validation study was exclusively collected and analysed for this paper.

### Experimental procedure

Facial expression production task. Our protocol was based on our previous work ^14^ which has shown to yield reliable communicative emotional signals when considering both signal dynamics and perceptual inference of emotion. In two separate sessions, across two days, participants’ facial movements were recorded during a facial expression production task. Participants were asked to signal three emotions (anger, happiness, and sadness) as they would naturally use their everyday interactions. This was done in two counterbalanced conditions. An *Expression only* condition where participants simply produced non-verbal facial expressions to signal each emotion, In the *Emotive speech* condition where, they were asked to express the same emotions while verbalising a “neutral” sentence (“Hi, my name is Jo, and I am a scientist”). This ensured that facial dynamic differences across emotions weren’t necessarily driven by the actual spoken content, but rather reflective of the target emotion dynamics. Similar approaches have been also used in the literature to generate datasets to study emotional production and perception stimuli with both linguistic and pseudo-linguistic (nonsense) expressive utterances ^43^. The conditions are motivated by the fact that in social interaction, facial expressions can be signalled both in isolation and embedded within *Emotive speech*, meaning that facial movements might signal information relevant to emotion and speech simultaneously, yet how speech and emotional cues are layered in facial dynamics is achieved is unclear. We focused on those expressions because they represent emotion signals which are often expressed in social interaction and communication ^58^. While we focus on three emotions only, the inclusion of a wider range of participants, speech and non-speech conditions, and data across multiple days provides richer representations of facial dynamics and variability compared to previous work in this area, which is often restricted to a few signallers, emotions, and conditions. Facial movements were recorded with participants seated with their head at the centre of a rectangular frame to ensure central positioning towards the camera, positioned 1 meter from a tripod-mounted video camcorder, sampling at 60Hz f/s (Sony Handycam HDR-CX240E). The start and end points were signalled using an audible beep, which allowed us to custom-align the recordings of facial expressions - see the schematic illustration in Fig 8. The *Expression only* videos had consistent duration of around 10 seconds, with some videos in the *Emotive speech* condition being slightly shorter (see analytical approach section below for time matching and alignment procedures).

**Figure 8.**
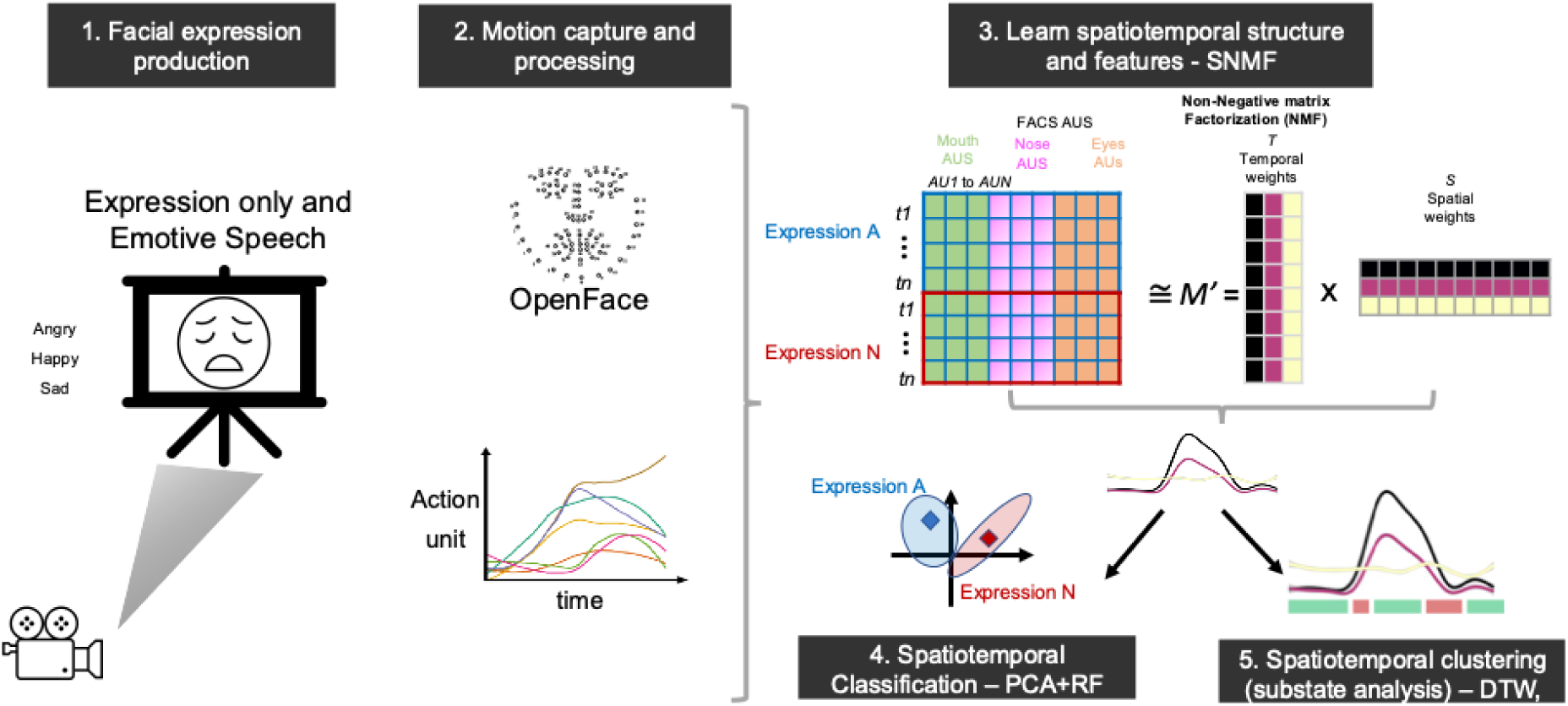
Schematic illustration of methodological and analytical approach. 1. illustrates the facial expression production task, where participants facial expressions (angry, happy sad) were recorded in two conditions (*Expression only* and *Emotive speech*). Expressions were perceptually validated in separate study and sample. 2. OpenFace was used to quantify AU patterns over time and align motion patterns. 3. summarises the procedure for the concatenation of AU timeseries and the application of Non-Negative Matrix Factorization (NMF) to learn an underlying low-dimensional spatiotemporal representation of the data. 4. The accuracy and effectiveness of this lower-dimensional representation were validated through a series of sensitivity and classification analysis. These relied on extracting timeseries features (e.g. curvature, complexity) from spatiotemporal components and using PCA to summarise them into a score per video, used for training the random forest classification model validated in an unseen set. 5. Time series clustering was used to identify substates or ’phases’ in facial movement components.

A separate perceptual validation study was conducted online with participants naive to the facial expression production task. Building on our validated protocol^14^, we used point-light recreations of the stimuli to isolate only spatiotemporal facial expression dynamics and explore correspondence with human observers’’ ratings and categorisation of emotion signals (see SI- Perceptual Validation Study)

### Analytical approach

#### Data processing

Facial expression production videos were processed using OpenFace ^59^, an open-source suite for automated face-tracking, landmarking and facial action unit detection based on the facial action coding system (FACS) ^2^. We focused on Action Units (AUs), which code for the observable presence and intensity of basic face actions (e.g., blinking, mouth widening, eyebrow-raising) that can be used to produce almost any facial expression movement. We estimated the temporal activation weights for 18 AUs. Only videos with a tracking accuracy of over 90% and with participant data across two sessions were used, resulting in a total of 532 videos with approximately 240 time points each. Based on visual inspection of the movement profiles, we applied a moving average (window = 3 frames) to smooth out the slight jitter inherent in face tracking and AU estimation, while preserving the underlying trends and dynamic range. To ensure an equal temporal window for spatiotemporal analyses, AU timeseries were reduced into 100-time bins for each video. Binning was preferred over more computationally expensive alternatives such as dynamic time-warping (DTW) for practicality since the overall timeseries duration was very similar across videos, and visual comparisons between binning and DTW tests on our data showed negligible differences – see Fig S11 in SI.

#### Spatiotemporal analysis

Timeseries of AU weights for the full facial expression production task were analysed using Non-Negative Matrix Factorization (NMF) ^60^. NMF is a multivariate dimensionality reduction method that decomposes a high-dimensional dataset into two non-negative matrices that form a part-based representation of the data (i.e., the original data is achieved by combining/adding components). The non-negativity constraint is particularly well suited for representing movement and AU data, which is typically positive, and the two-matrix part-based decomposition can intuitively represent spatial vs time-varying aspects in the data – see Fig 8 (see also: ^22,40,60^).

The NMF takes the concatenated AU timeseries as input, denoted by a matrix ***M***, which is a product of two low-dimensional matrices (***T*** and ***S***), ***T*** represents the temporal weights (basis values for time bins), and ***S*** represents the spatial weights (face action activation weights). The product of ***T*** *and **S*** matrices denoted by ***M’*** represents an approximation of the full original matrix such that ***M∼M’ = T * S*.** The algorithm works by initialising the values for ***T*** and ***S*** (e.g., with random values) and using an iterative multiplicative update rule to sequentially update *T* and *S* by a factor that minimises the approximation error (i.e., residuals). While the initialisation and iterative nature of NMF means that the solution is non-deterministic (i.e., you can learn multiple equally effective part-based representations), these update rules have been shown to guarantee convergence (i.e., stability of the cost function) after a relatively low number of iterations ^61^. The optimal number of spatiotemporal components (*k*) was assessed computationally by fitting a range of values (*k* = 2 to 6) and choosing a *k* that optimised various model fit and reconstruction metrics (e.g., residual sum of squares, silhouette for clustering cohesion, cophenetic coefficient, see below). We also evaluated the final solution against NMF results on a permutated dataset, using block-based time shuffling. This resulted in an appropriate null dataset that destroyed the inherent spatiotemporal pattern in the data but retained some temporal ordering. We then compared fit metrics between original and permutated NMF (see also *Classification and clustering analysis*). We used the NMF decomposition functions in the NMF package for R ^62^. To visualise our NMF results in an intuitive manner, we trained a custom model to predict aligned Facial Landmark displacements (through Procrustes analysis) from AUs using partial least squares regression. Note however that the AU to Landmark face model is not a perfect one to one map between NMF results and visualisation, given the inherent many-to-one relationship between facial landmarks and AUs ^51^. For clarity of presentation and interpretation, we report the NMF components fitted separately for the *Expression only* and *Emotive Speech conditions*. However, sensitivity analyses confirm that these results are consistent with those obtained from a joint analysis of both conditions, thereby showing reliability of the identified spatiotemporal patterns, and that the results are not arbitrarily biased by the input data.

#### Multivariate timeseries classification

We used a supervised classification approach to validate the diagnostic value of spatiotemporal components for emotion expression. This involved five steps: (1) **Data splitting**: We began by splitting the original pre-NMF data into training (80%) and testing (20%) sets, ensuring that each set contained complete AU time series for each video. This resulted in a matrix with approximately 183 videos x 100 time points and 18 AUs for train, and 45 x 100 x 18 for test set, where emotions were all equally balanced (61 per category: angry, sad, and happy for train, and 15 each for testing).

This split was performed before any further analyses, and all subsequent steps maintained this separation ensuring further processing and model training was only done on the training set and then applying results to the test set ^63,64^. Hence any mention of “training”, always refers to operations on the respective training set. (2) **NMF re-training and projection:** We then re-trained the NMF model using only the training set data and projected the test set onto the trained NMF model space to ensure consistency of components for classification.

(3) **Feature extraction:** Because standard classification approaches do not account for inherent spatiotemporal features of the NMF components, we opted for an approach that represents the spatiotemporal structure via interpretable theoretic time series features (e.g., curvature, complexity) in line with best practices and established computational frameworks for multivariate timeseries features extraction classification ^65,66^. This led to a total of 40 features extracted per component. However because it is impractical to detail the individual computation of every single feature, for completeness we provide a full list of extracted features with a brief description in the Supplementary Information (Table S5).

The focus on timeseries features is motivated by previous research showing the role of timeseries features in capturing diagnostic dynamics for classification of facial expressions and body movements ^11,14,67,68^.

Given the large dimensionality of the data, a Principal Component Analysis (PCA) was used to summarise the timeseries features based on variance in the train set while retaining a number of dimensions equal to the number of NMF components for consistency. The test set features were then transformed using the PCA model fitted on the training set features to be used in the next step. The matrix provided to PCA mirrors the number of videos from the original NMF analysis and the total number of extracted spatiotemporal features per component. Specifically, for the training sets under each condition (*Expression only* and *Emotive speech*), the matrix was 183 videos by 114 features (38 features for each of the three spatiotemporal components), while the test sets are 45 by 114. The component scores were used as inputs for classification analysis in the next step. Conceptually, each principal component (PC) captures the temporal dynamics inherent in the original NMF components, effectively consolidating the complex features patterns into a single feature score per spatiotemporal dimension. On average, the first three PCs account for between 34% and 40% of the variance in the data across both the *Expression only* and *Emotive speech* conditions, in both the training and test sets, showing that they capture structure without overfitting the facial behaviour data.

(4) **Classification training and testing.** We then trained a classifier on spatiotemporal component features derived from step 3 and applied to predict emotion expression in the held-out test set. Specifically, we used a Random Forests (RF) Classifier - a supervised statistical learning method that learns a discriminant model between a set of input variables (e.g., spatiotemporal component features) and a set of target classes (expressions). Note, that we achieved similar results with different methods (e.g., Support Vector Machine, Linear Discriminant Analysis), thus our results are not a function of the analytical approach. The Random Forests Classifier has the advantage of being relatively more robust to overfitting and able to capture non-linear relationships and therefore was preferred. We used cross-validation during training, where multiple training sets are created, and the data not included in the test set is used to test the accuracy and tune the model parameters. Reliable classification on the test set would indicate that the underlying spatiotemporal components (identified by the NMF analysis) carry diagnostic information for differentiating expressions. Note that the theoretical chance level for our three-way classification problem (i.e. happy, sad, and angry expressions) is defined as 33%, and because the datasets are balanced, this is also the empirical “no-information rate” or null rate. In addition, we derived significance tests for our classification results using binomial tests to determine whether the model’s accuracy on the test set was significantly greater than what would be expected by random guessing (by comparing it to the no information rate). These binomial p-values were compared with permutation-based p-values, which were generated by comparing the observed accuracy to the distribution of accuracies obtained from shuffling the labels (expressions) in the test set (N = 1000). The negligible differences between the binomial and permutation-based p-values for our dataset validated the robustness of our results, so we primarily report the binomial p-values.

As an additional sensitivity test, we used Fisher’s Exact Test to assess the statistical significance of the confusion matrix, by comparing pairwise expression differentiation with adjustments for multiple comparisons. The null hypothesis is that there is no association between rows and columns, or in other words differences in cell values (frequency) are due to chance. A low p-value in Fisher’s exact test indicates statistically significant differences between pairs, meaning the pattern in the confusion matrix is unlikely to have occurred by chance.

While a handful of previous studies previous work has employed elements of our proposed pipeline such as the dimensionality reduction of facial movement data across entire datasets, our approach is unique in that it explicitly builds in the generalisability and robustness of the spatiotemporal patterns we uncover. Furthermore, the spatiotemporal classification pipeline enables us to directly link the inherent dynamics (represented through the extracted features) to their diagnostic value in differentiating emotions. We also used the same general approach above (from step 4) to analyse the perceptual validation data, linking our spatiotemporal components and human observers’ perceptual emotion ratings, for the held-out test set – see ^69,70,71^. Full details are provided in the SI - Perceptual validation of the spatiotemporal components for emotion signalling.

#### Spatiotemporal clustering for substates analyses

We investigated whether substates could be extracted from the spatiotemporal patterns of *Expression only* and *Emotive speech* facial movements. We considered substates as local intervals where the components exhibit salient signatures that are consistent across expression timeseries. We operationalised this as a segmentation of a multivariate timeseries into finer temporal segments with arbitrary lengths. To do this we first derived speed and displacement for spatiotemporal components (from step 2 above - NMF re-training and projection), as these features are sensitive to segments local characteristics of other body movements (e.g., swing and stance signatures in walking, fixation and saccades in eye movements) ^23,34,35,72^. These were computed using standard approaches from the kinematic literature. Speed was computed as the first derivative of the time series, approximated by the bin-wise difference between consecutive values and normalised by the corresponding time interval. Displacement was derived as the differencing of the timeseries over time relative to the starting values (baseline).

Second, we applied spatiotemporal clustering using the *dtwclust* package in R, using the component speed and displacement metrics as inputs. This was motivated by the fact that, unlike other movements with well-established kinematic profiles, we have no *a priori* knowledge regarding the characteristics of facial expression substates. Given the computational complexity of DTW, we first tested cluster sizes on a smaller subset of *Expression only and Emotive speech* data. We used exploratory visualisations (e.g. Fig 6) which suggested two to three interpretable segments described transition and stability phases in spatiotemporal components of facial expressions in line with the hypothesised patterns of transition, sustainment and relaxation of AUS. Note that because the substate clustering inputs are based on spatiotemporal components representing AUs, and because AUs are equally valid descriptors of facial behaviour in both non-verbal expression and emotive speech, substate analysis are equally valid for both conditions. Therefore, any differences in substate patterns are reflective of condition-specific nuances, rather than a bias to specific condition.

Because the clustering results are mostly descriptive and return only cluster labels for partitioning the input data, it does not inherently provide a statistical characterisation for why these clusters emerge. To facilitate intuitive interpretation of clusters, we therefore visualise the pattern of clusters and the input data (Fig 6). For example, it became apparent that clusters had distinct speed signatures but also apparent sequences that are more prominent in *Expression only* and more dynamic in *Emotive speech* conditions. To statistically assess these patterns, and explore how they differentiate emotion and conditions, we computed two metrics: (1) speed within the substate and (2) entropy; both which have been shown to capture diagnostic information about facial expression movement patterns ^11,14^. Specifically given that speed contributes to the clustering as an input, statistically comparing speed gives direct insights into how the clusters were generated (see Fig 6). Entropy, is on the other hand is an information-theoretic measure of complexity, directly related the dynamics of state transitions (or sequences) within a system ^44^. In this context, entropy provides a suitable summary of the dynamics of substates that can be computed post hoc (after cluster generation) and summarised in a single statistic, describing how complex or unpredictable the sequence of these substates is (e.g., how much information about the current substate can predict the next substate). This statistic aimed at exploring the apparent differences in the temporal patterns of substates between expressions and conditions (see Fig 6). We computed entropy using the transition entropy function (for state change entropy) from the *GrpString* package for R ^73^ but for simplicity we refer it as entropy. To improve interpretability, entropy was normalised by dividing it by the maximum theoretical entropy for a system with the same number of substates.

Both speed and entropy measures were computed for each individual expression and analysed using a mixed-effects model to test the effect of emotion (happy, angry, sad), condition (*Expression only*, *Emotive speech*), and substate (relaxed, transition, sustain) as fixed effects and subject (face id) as a random effect (random intercept). Fixed effects terms were also initially included as random effects (slope terms for expression and condition), however, random structure was simplified in line with best practice for parsimonious mixed models ^74,75^ which revealed that “maximal models” (full complex slope models) tended to overfit (according to AIC and BIC indices) and lead to instability in convergence.

As such the final models had the form:

speed (or entropy) ∼ emotion × condition × substate + (1 + condition | face ID)

We used a linear mixed models as opposed to the classification pipeline described earlier given the primary aim to characterise substates and statistically assess differences between emotion and conditions on the basis of substate patterns. Additionally, the more complex and nested structure of the substate data makes classification overly complex to structure and present. Linear mixed models were fitted using the lme4 package for R ^76^ with inferential tests derived from lmerTest^77^ using the Satterthwaite’s approximations for degrees of freedom from the covariance structure and report Type III tests of fixed effects, in line with best practice recommendations ^74,75^. Pairwise and interactions post-hoc were all adjusted for multiple comparison using Bonferroni corrections.

### Perceptual validation experiment

#### Participants

An additional sample of 45 naïve participants (22 female, age range: 23-40), took part in an online study to validate the stimuli from the facial expression production study described in the main manuscript. Participants were recruited via Prolific (an online testing platform) and were compensated at £9 per hour. Participants were selected to have English as their first language and residing in the UK, with no current mental health conditions or autism diagnosis ^78^.

#### Stimuli and perceptual validation task

Given the online format and the large number of stimuli derived from the facial expression production task, it was not feasible for participants to rate every single video. 90 selected videos from the spatiotemporal analyses (held-out test-set), were used as a representative subset of the stimuli in the study (45 for *Expression only* and 45 from the *Emotive speech* conditions, with 15 videos per emotion: happy, angry, and sad). To isolate the motion characteristics of the original stimuli, we used face landmarks from openFace (see Methods)^79^ to generate the stimuli for the perceptual validation. Specifically, we used the aligned facial landmarks to generate point-light face display (PLFDs) animations, building on a validated protocol used in our previous work ^14^. To correct for inherent jitter in facial landmarking, a slight smoothing was applied to facial landmarks using a moving average with a window of 2 frames.

#### Task

To ensure attentiveness and authenticity of participants, we implemented random video checks throughout the task using their webcam (with their consent) to capture snapshots of participants while they completed the task, in addition to a self-report survey in the end of the experiment. Each video was displayed for its original duration (approximately 10 seconds). For each video, participants rated the intensity of three emotions (anger, happy, and sad) on a scale from 0 to 100. An additional “other emotion” rating was included to ensure variability and complexity, thus reducing the likelihood that participants would simply eliminate options rather than provide genuine ratings. To reduce parity in the different ratings, participants were instructed to assign their highest rating to the emotion they perceived as most dominant in each video. The presentation order for each condition (*Expression only* and *Emotive speech*) was counterbalanced across participants and videos were presented randomly. On average, the entire task took about 35 minutes to complete.

#### Perceptual validation of facial expression production

For each video, and participant, we assigned the perceived emotion categorisation based on the highest rating given by the participant. For instance, if the highest rating was 70 for “happy” and lower for all other options, that trial was categorised as “happy”. This categorisation was performed both on the full dataset and on the aggregated dataset where ratings were averaged per stimulus across participants. To assess the alignment between perceptual ratings for emotion and the facial expression production categories, we tested for the correspondence between perceptual derived emotion categorisations, and the intended “ground-truth” emotion from the facial production task. This was tested using Pearson’s Chi-squared tests - conducted with 2000 simulated replicates. A significant test, indicate that the perceptually derived categories closely reflect the production-based “ground-truth”, thereby providing strong validation for the stimuli.

#### Correspondence between spatiotemporal models and human observers

Building on the data described above, in study 2, we conducted two additional analyses to validate the predictive value of spatiotemporal components to perceptual processing of emotion from facial signal dynamics. First, we conducted a correspondence analysis to test the relationship between emotion classification predictions derived from our spatiotemporal-based approach and those provided by human observers on the held-out test set. The use of held-out data (not used in the training of the NMF analysis) inherently provides a generalisability test, ensuring that any correspondence is not simply driven by overfitting to characteristics that differentiate the expressions in the data.

Second, similar to the use of the spatiotemporal components to predict emotion categories in the held-out test set from production only data (see Multivariate timeseries classification, step 4 in the manuscript), we used the spatiotemporal components to directly predict human emotion categorisation. This complementary analysis allows us to test not only whether our model’s predictions align with participants’ ratings, but also whether the spatiotemporal features themselves are directly predictive of perceptual judgments. Both approaches described above align with the current practice in linking model-based performance with human behaviour in neuroscience and machine learning, which has been used in modelling human visual processes ^69,70^ – although see ^71^.

For consistency we use the similar classification approach described in the spatiotemporal classification section, relying on a Random Forest classifier. However, to ensure reliability and generalisability of these findings, we employed two validation approaches: (a) Leave-One-Stimulus-Out (LOSO): For the data averaged across participants, the model was iteratively trained on all data except for one stimulus at a time. This procedure ensures that prediction of human observers’ emotion categorisations generalise across different stimuli; (b) Leave-One-Participant-Out (LOPO): In this approach, the model was trained on data from all participants except one and then tested on the held-out participant. This process was repeated until every participant had served as both training and test data, confirming that the results are robust to variability in individual participants ratings.

High accuracy scores indicate that the spatiotemporal components robustly predict perceptual ratings. Although the theoretical chance level for a three-way classification (angry, happy, sad) is 33%, the empirical chance level may be higher if participant responses are skewed toward a majority category (e.g., happy). In such cases, using the majority response as the baseline provides a more stringent and data-specific test of model performance.

## Data availability statement

Supporting data is available in the Open Science Framework (OSF): https://osf.io/3nzk7/?view_only=db7d9467423f42b794385e549b6aa96a All code for pre-processing and models is available at https://github.com/hcuve/spat-tmp-emo-faces

## Acknowledgments

We would like to thank Professor Rachael Jack for her insights during the initial discussions on facial signalling dynamics and dimensionality reduction. We would also like to thank Katherine Lane and Tahniyyat Bokhari, for their assistance with data collection for a portion of the study.

## Author Contributions (CRediT)

HCC, Conceptualisation, Data Curation, Formal Analysis, Funding Acquisition, Investigation, Methodology, Software, Supervision, Validation, Visualisation, Writing – Original Draft Preparation, Writing – Review & Editing, Resources SSC, Methodology, Validation, Writing – Review & Editing JC, Conceptualization, Funding Acquisition, Methodology, Writing – Review & Editing, Resources

