## Supplementary Info for "Spatiotemporal dynamics and substates underlie emotional signalling in facial movements"

Hélio Clemente José Cuve[1*], Sophie Sowden-Carvalho[2], Jennifer Cook[2]
[1]School of Psychology and Neuroscience, University of Bristol, Bristol, United Kingdom
[2]School of Psychology, Centre for Human Brain Health, University of Birmingham, Birmingham, United Kingdom

### Perceptual Validation of Facial Expression Production

There was significant correspondence between the perceptually derived emotion categories and the intended production-based classifications for both *Expression only* ($X^2 = 36.922$ $p < .001$) and *Emotive speech* ($X^2 = 20.28$ $p < .001$). These indicate that the emotion ratings provided by participants closely mirror the facial production categories, thereby confirming that the stimuli reliably evoke the intended emotional signals. This can also be visualised in Figure S1, where in general, the dominant emotion rating slider aligns with the "ground-truth" expression category. In line with previous research, the rating patterns in Fig S1 also suggest that happy expressions are the most distinctive perceptually [1,2].

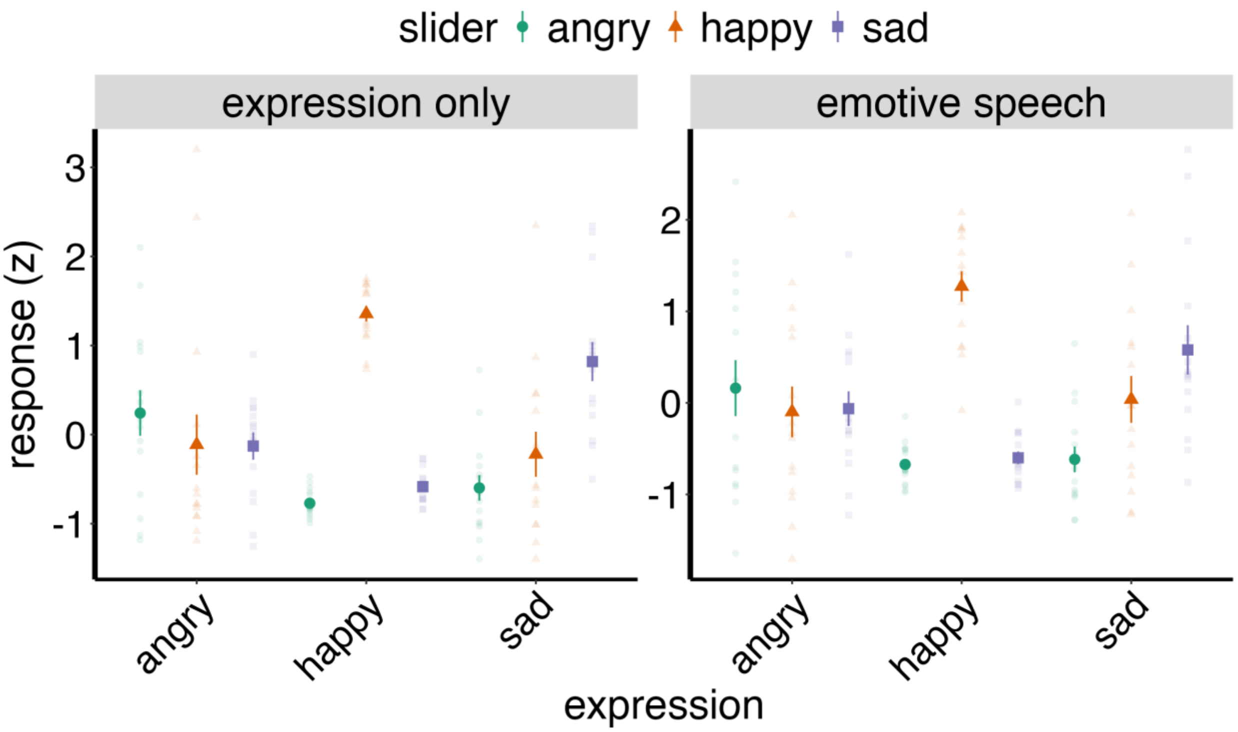


Figure S1. Perceptual ratings validation of facial expressions.

Note. Mean and individual standardised (z-scored) emotion ratings for angry, happy, and sad expressions in the *Expression only* (left) and *Emotive speech* (right) conditions. Each point represents the average rating for a single video (aggregated across participants) for that emotion. The perceived emotion ratings (response, y-axis) were converted to z-scores for visualisation. The dominant perceptual emotion rating aligns with the intended facial expression production emotion “ground-truth” (x-axis), confirming that the stimuli reliably evoke the intended emotional signal.

### Expression only condition - NMF validation analysis

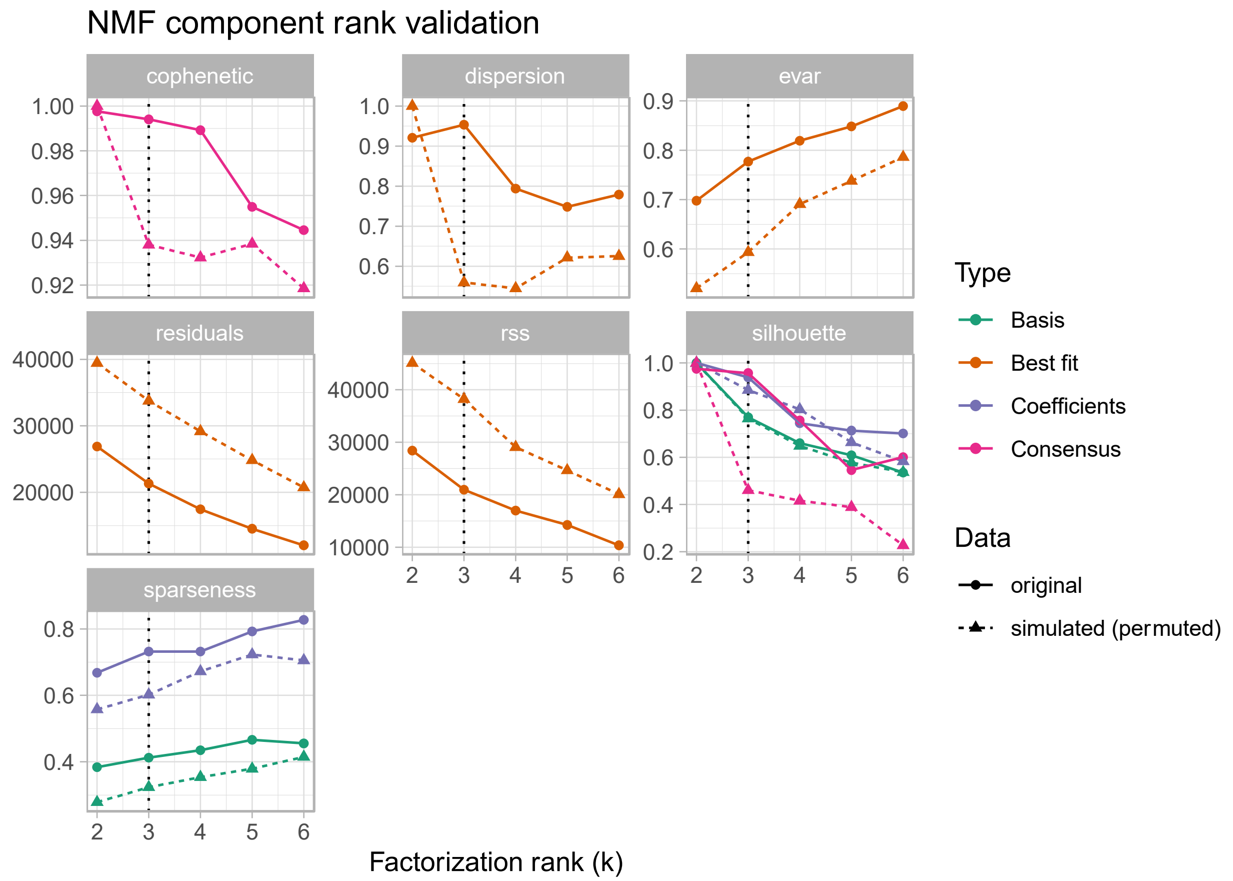


Figure S2. Validation analysis comparing NMF on real expression only data against permuted data.

To validate the NMF model and determine the optimal number of components, we compared various fit measures obtained from the original data against those derived from NMF applied to permuted data, which disrupts the inherent spatiotemporal patterns. The NMF model fitted on the original data consistently shows a better fit than the random data NMF across several metrics (e.g. higher silhouette scores and lower residuals), particularly favoring a three-component solution. This contrast highlights the model's effectiveness in capturing meaningful patterns in the original data compared to the noise-driven results from the permuted data.

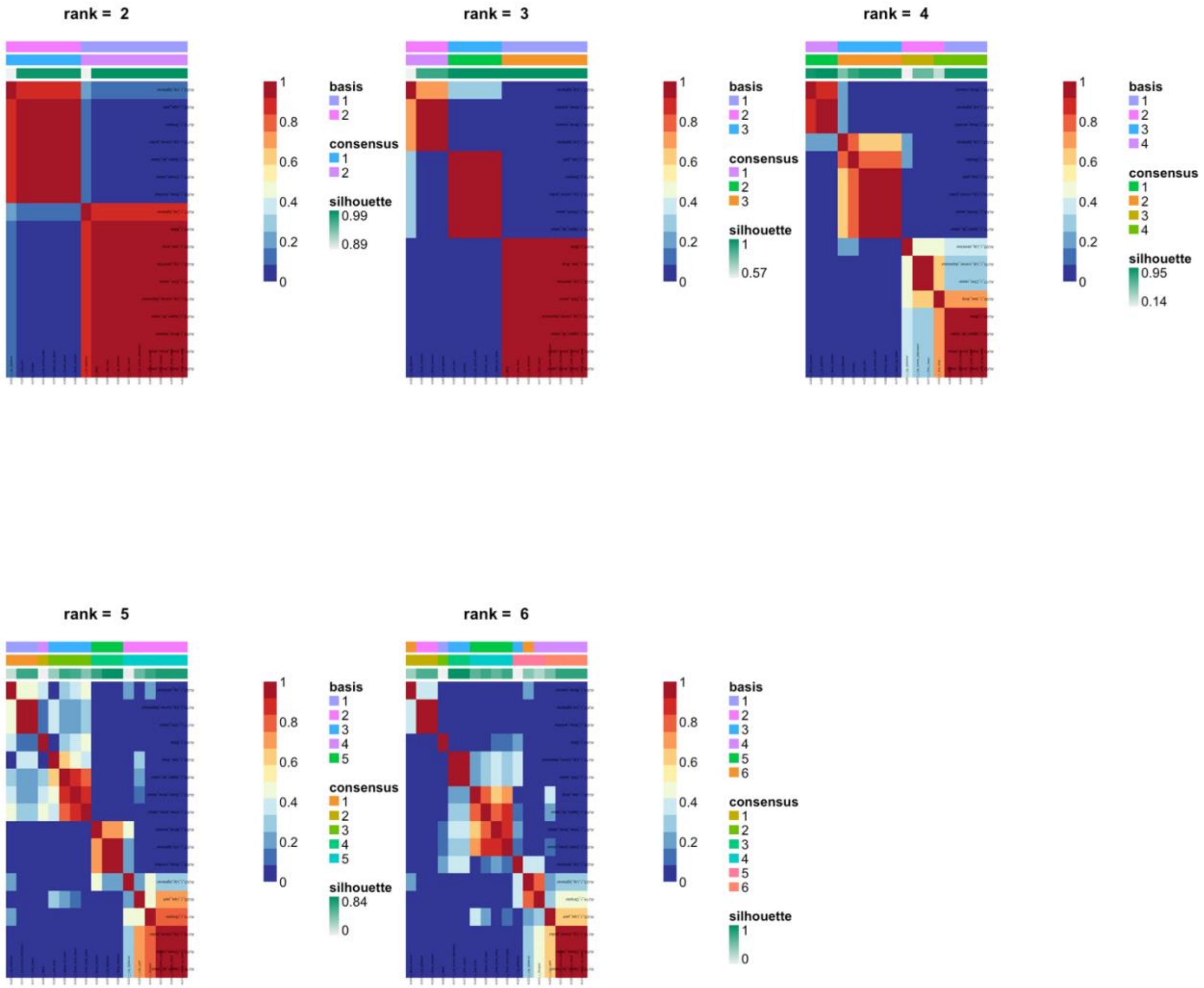


Figure S3. Consensus map for expression-only NMF across a range of k values. The consensus map illustrates how consistently pairs of data points are assigned to the same component clusters across multiple NMF runs with different initial conditions. The grid-like diagonal patterns with high silhouette values indicate stable clustering solutions. Specifically, this map suggests that 2 to 3 components provide the most consistent (stable) clustering, as evidenced by the high silhouette scores and strong consensus. Beyond k = 3, particularly at k = 4 and 5, the stability of the clustering significantly degrades, indicating that these higher k values do not capture the data's structure as reliably.

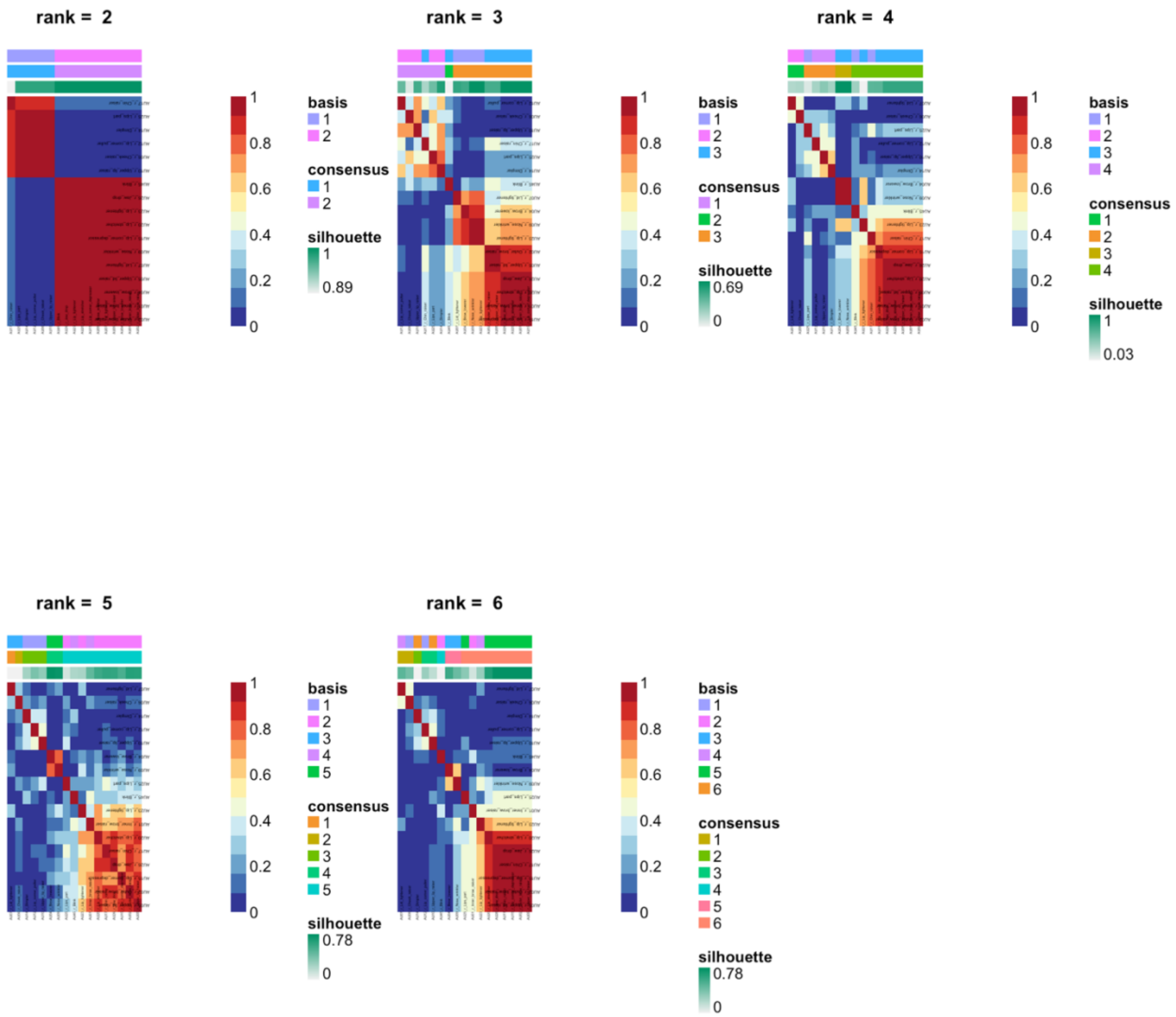


Figure S4. Consensus map for NMF on permuted expression-only data across a range of k values

Consensus results for NMF applied to permuted expression-only data across a range of k values. Unlike the original data, the permuted data lacks inherent spatiotemporal structure, and this is reflected in the consensus map. The absence of well-defined diagonal patterns and generally low consensus values indicate that the NMF model struggles to consistently assign data points to the same component clusters across different runs. This instability, especially as k increases, underscores that the meaningful patterns identified in the original data are disrupted when the data is permuted. While at k = 2 the model is stable, combined with fit metrics, suggest that it is overfitting noise.

**Expression only condition- classification evaluation metrics**

| | Reference | | |
|---|---|---|---|
| Prediction | angry | happy | sad |
| angry | 14 | 0 | 3 |
| happy | 1 | 15 | 0 |
| sad | 0 | 0 | 12 |

Table S1. Confusion table for expression only classification results. Metrics are calculated on the held-out the test-set.

| | Class: angry | Class: happy | Class: sad |
|---|---|---|---|
| Sensitivity | 0.93 | 1 | 0.8 |
| Specificity | 0.9 | 0.97 | 1 |
| Pos Pred Value | 0.82 | 0.94 | 1 |
| Neg Pred Value | 0.96 | 1 | 0.91 |
| Precision | 0.82 | 0.94 | 1 |
| Recall | 0.93 | 1 | 0.8 |
| F1 | 0.87 | 0.97 | 0.89 |
| Prevalence | 0.33 | 0.33 | 0.33 |
| Detection Rate | 0.31 | 0.33 | 0.27 |
| Detection Prevalence | 0.38 | 0.36 | 0.27 |
| Balanced Accuracy | 0.92 | 0.98 | 0.9 |

Table S2. Evaluation metrics by expression – expression condition only. Chance is .33. Metrics are calculated on the held-out the test-set.

### Emotive speech condition- NMF validation analysis

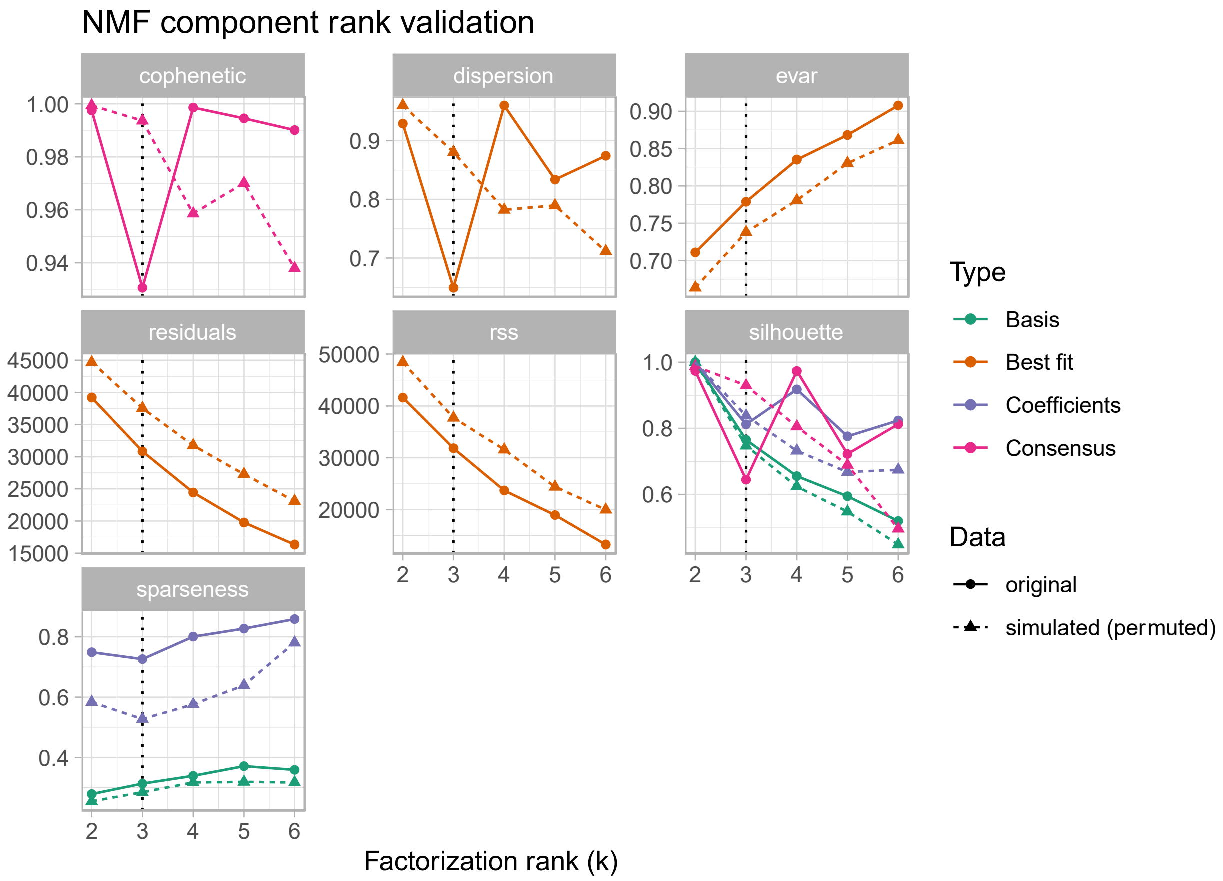


**Figure S5. Validation analysis comparing emotive speech NMF against NMF on permuted dataset**

Comparative validation analysis for NMF applied to emotive speech data versus permuted emotive speech data across a range of k values. The results show that the NMF model applied to the original emotive speech data consistently provides a better fit (across a range of metrics) compared to the permuted data (e.g. higher silhouette scores and lower residuals). This suggests that the NMF model is effectively capturing meaningful patterns in the emotive speech data. The fit metrics also highlight that the most stable and reliable component solution occurs around k = 3, where the model shows a balance between fit quality and stability.

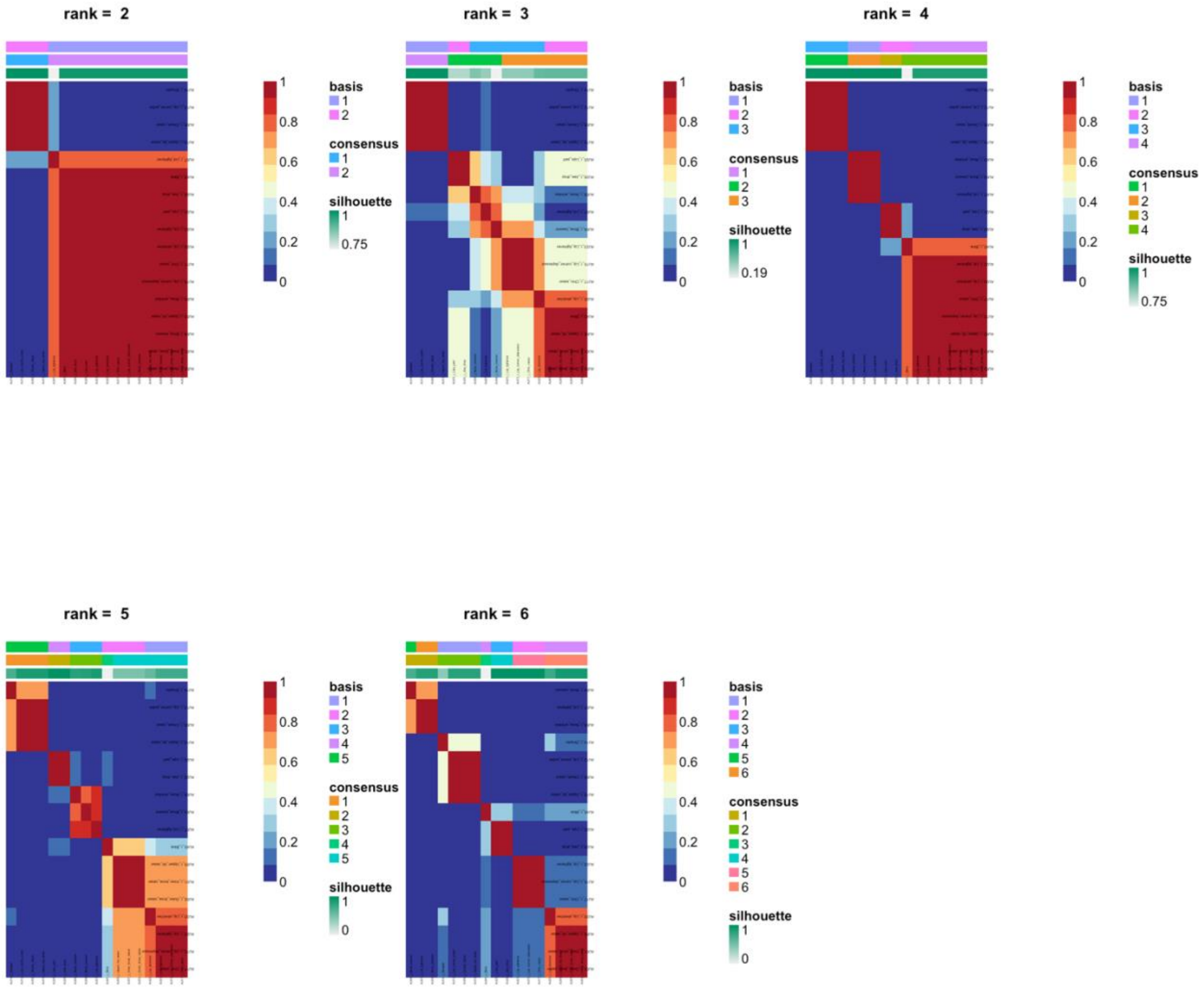


Figure S6. Consensus map for emotive speech NMF across k values
The presence of high consensus values, particularly at k = 3 to 4, suggests that the NMF model reliably and consistently groups data points into the same components, indicating that this k value is optimal for capturing the emotional structure within the speech data.

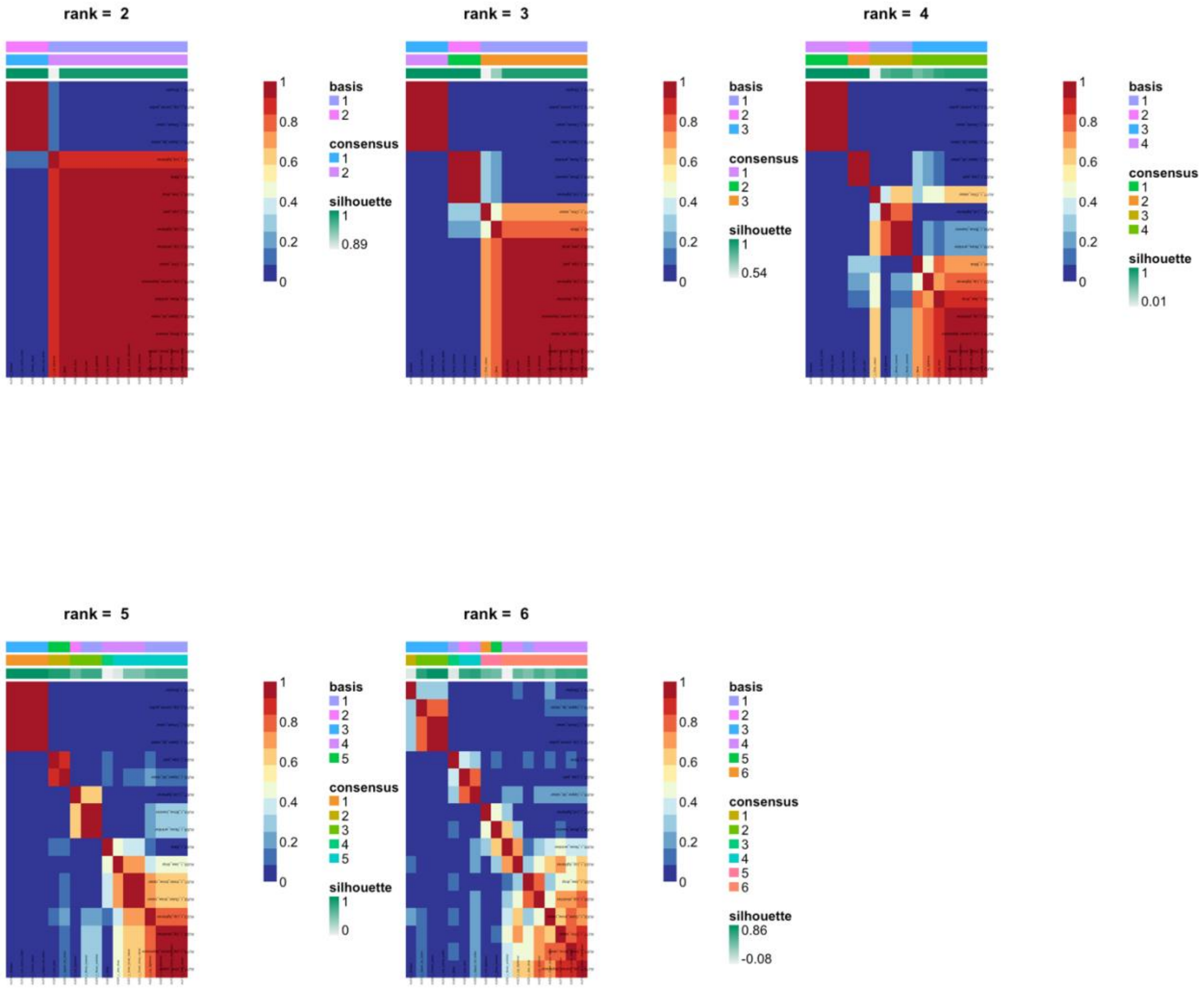


Figure S7. Consensus map for NMF on permuted emotive speech data.
While some stability might be observed at lower k values, taken together with the fit metrics, this suggest that NMF on random data is overfitting to noise rather than capturing meaningful clusters.

**Emotive speech classification evaluation metrics**

| | Reference | | |
|---|---|---|---|
| Prediction | angry | happy | sad |
| angry | 8 | 2 | 4 |
| happy | 0 | 13 | 1 |
| sad | 7 | 0 | 10 |

Table S3. Confusion table for emotive speech. Metrics are calculated on the held-out the test-set.

| | Class: angry | Class: happy | Class: sad |
|---|---|---|---|
| Sensitivity | 0.53 | 0.87 | 0.67 |
| Specificity | 0.8 | 0.97 | 0.77 |
| Pos Pred Value | 0.57 | 0.93 | 0.59 |
| Neg Pred Value | 0.77 | 0.94 | 0.82 |
| Precision | 0.57 | 0.93 | 0.59 |
| Recall | 0.53 | 0.87 | 0.67 |
| F1 | 0.55 | 0.9 | 0.62 |
| Prevalence | 0.33 | 0.33 | 0.33 |
| Detection Rate | 0.18 | 0.29 | 0.22 |
| Detection Prevalence | 0.31 | 0.31 | 0.38 |
| Balanced Accuracy | 0.67 | 0.92 | 0.72 |

Table S4. Evaluation metrics by expression for emotive speech condition. Chance is .33. Metrics are calculated on the held-out the test-set.

### Comparison of joint vs separate spatiotemporal analysis of *Expression only* and *Emotive speech*

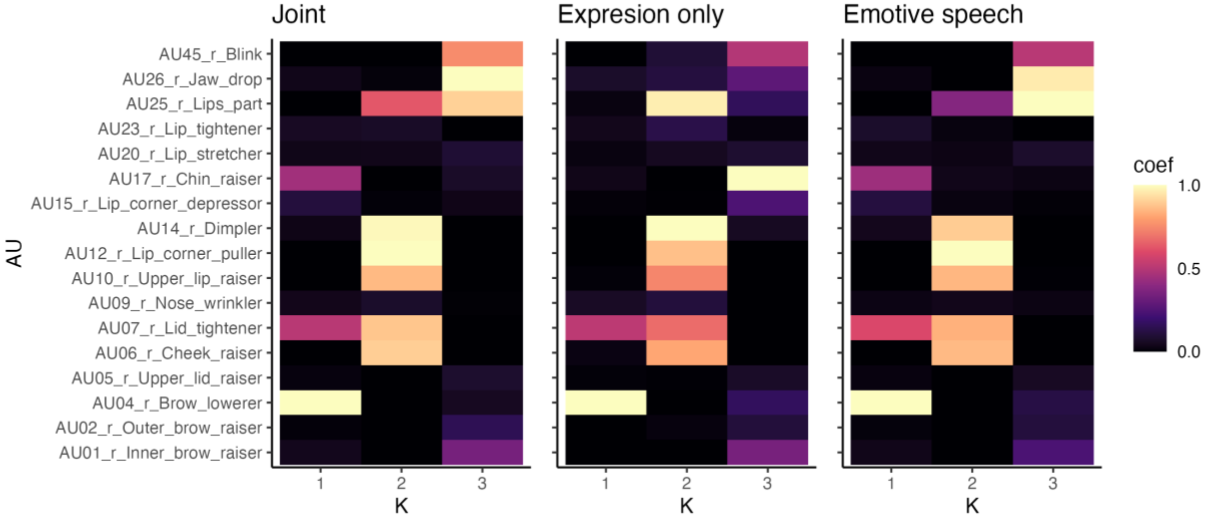


Figure S8. AU-component profiles for joint and individual spatiotemporal analysis for *Expression only* and *Emotive speech.*

Note. Heatmaps of NMF-derived Action Unit (AU)-component profiles for a joint analysis (left) versus separate analyses of the *Expression only* (centre) and *Emotive speech* (right) conditions. Warmer colours indicate stronger AU loadings for each component (K=1–3). *Coef.* = NMF coefficient. The similar AU–component patterns across the three panels confirms that most components are shared between conditions, demonstrating the robustness of the identified spatiotemporal patterns. Subtle visual differences reflect condition-specific nuances and small effects of facial landmark normalisation. For instance, in the *Emotive speech* condition, there is greater emphasis on lower-face AUs involved in speech articulation - such as AU25 (Lips part) and AU27 (Lip stretcher) - across Components 2 and 3. By contrast, the Expression Only condition shows these lip-related AUs more parsimoniously in a single component (2). The joint NMF plot preserves the overall pattern while effectively combining the condition-specific nuances.

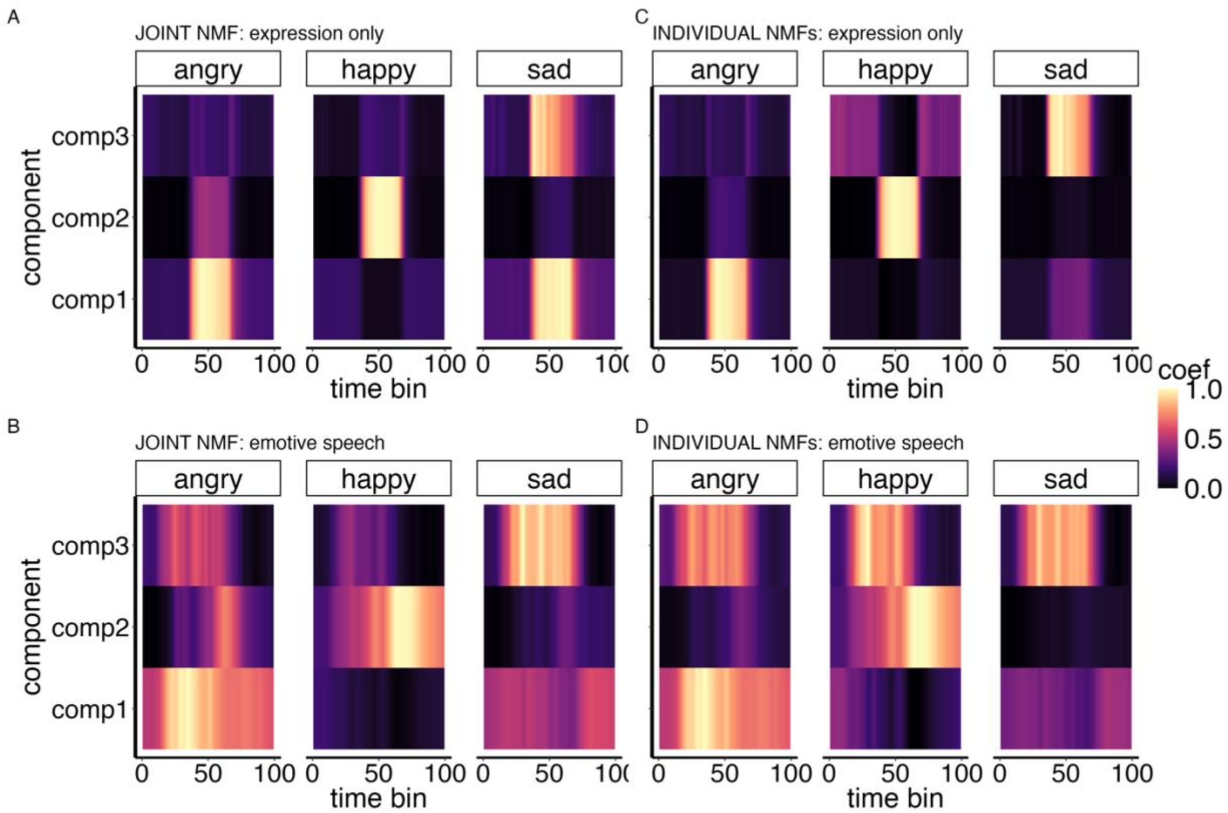


Figure S9. Spatiotemporal component profiles for joint vs individual spatiotemporal analysis for *Expression only* and *Emotive speech*

Notes. These plots compare the overall temporal evolution of spatiotemporal components derived via NMF when applied jointly (A and B) and individually (C and D) for each condition (*Expression only* and *Emotive speech*). The profiles remain largely consistent across conditions, demonstrating the robustness of our NMF approach. For example, the dominance of Component 1 in capturing the 'happy' expression is evident in both the joint and separate analyses. This shows that the spatiotemporal components are reliably and that any differences observed between conditions are reflective of genuine, condition-specific nuances rather than artefacts of the input signals or methodological instability.

### Perceptual validation of the spatiotemporal components for emotion signalling

#### Correspondence between spatiotemporal component-based emotion classification and human observers

There was a significant correspondence between the expression labels predicted by the spatiotemporal model on the held-out test set and the categorisations derived from participants ratings for both *Expression only* ($X^2 = 33.125$, $p < .001$), and for the *Emotive speech* conditions ($X^2 = 15.622$, $p = .002$). These results indicate that the spatiotemporal component-derived emotion classifications align well with human observer classifications. In other words, the spatiotemporal components capture meaningful patterns that reflect how humans perceive and interpret emotional signals from facial dynamics.

This finding is further illustrated in Figure S10, which shows that the three expressions (angry, happy, sad) remain broadly differentiated based on spatiotemporal model predictions. Notably, happy expressions are more strongly differentiated overall, whereas angry expressions appear less distinct in the *Emotive speech* condition. The latter may be partly attributed to the inclusion of the "other emotion" rating scale, which was deliberately introduced to increase variability. Nonetheless, results support that the low-dimensional spatiotemporal components effectively captures emotional signals in a manner that is consistent with human perception. The use of held out set and an independent sample, underlie the robustness and predictive value of the spatiotemporal components.

#### Predicting human emotion categorisation of dynamic facial signals directly from spatiotemporal components

While the analysis above shows correspondence of spatiotemporal-based emotion classification and participants perceptual categorisations, it doesn't directly relate the spatiotemporal components underlying to participants categorisations. Consistent with the previous results, the low-dimensional spatiotemporal components of facial expressions were predictive of participants' emotion categorisations of PLFDs. In the *Expression only* condition, spatiotemporal components accurately predicted participants' categorisations both when aggregated by stimulus (ACC: 0.6778, 95% CI: [0.571, 0.7725], $p < 0.001$) and across participants, using a leave-one-participant-out cross-validation approach (ACC: 0.66, 95% CI: [0.641, 0.685], $p < .001$). Similarly, the spatiotemporal components learned from facial production data predicted perceptual categorisations of *Emotive speech* across stimuli moderately well (Accuracy: 0.6889, 95% CI: [0.5335, 0.8183], $p = 0.08$), and strongly across participants (ACC: 0.6072, 95% CI: [0.584, 0.629], $p < .001$) - (see also Figure S-11).

In summary, these findings indicate that participants' ratings of emotion in point-light versions of the stimuli - preserving the spatiotemporal dynamics - are robustly captured by the low-dimensional spatiotemporal structure learned from production data. This supports our claim that the low-dimensional spatiotemporal structure is not merely an artefact of dimensionality reduction but rather reflects functionally and perceptually relevant information about emotional facial expression.

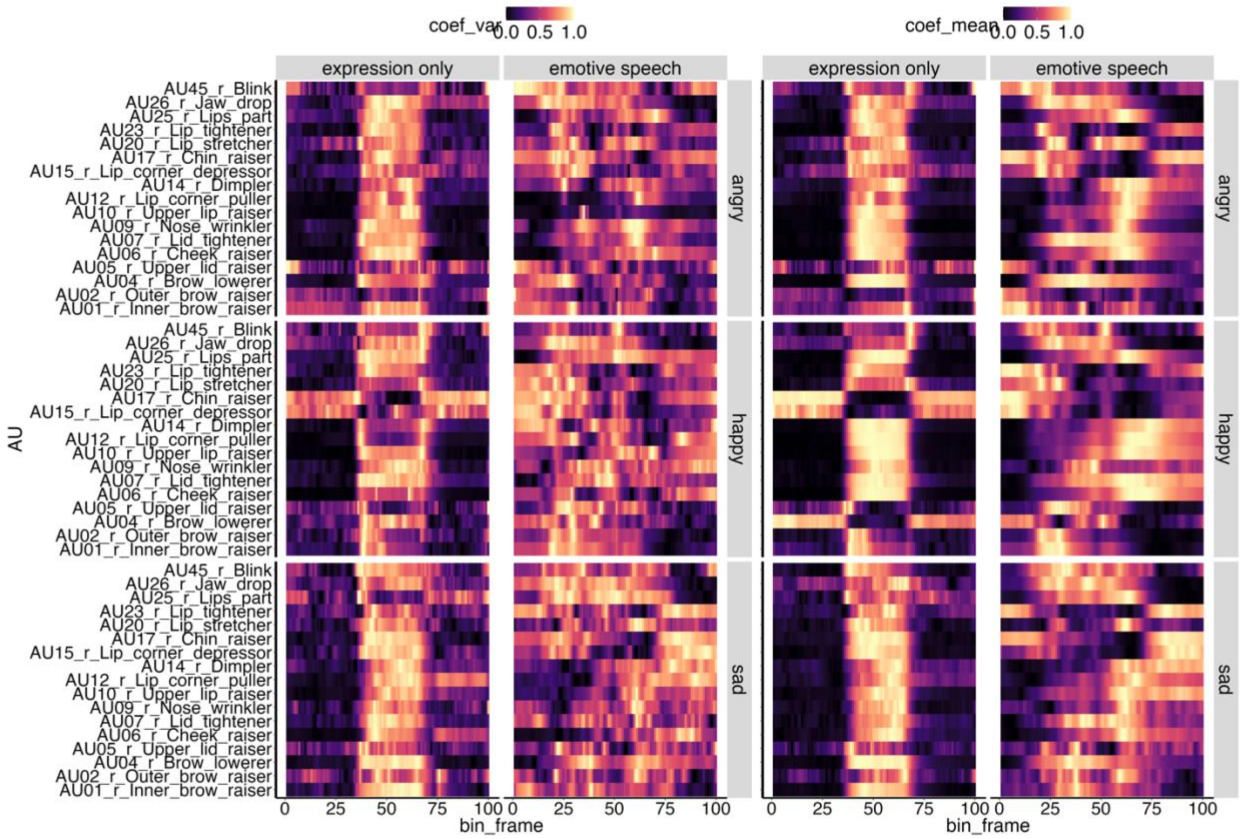


**Figure S10.** Heatmaps illustrating the distribution of facial Action Units (AUs) over time for *Expression only* (left panels) and *Emotive Speech* (right panels) conditions and each target emotion (*angry*, *happy*, and *sad*). Warmer shades (yellow/orange) indicate higher mean or variability (standard deviation) while cooler shades indicate lower values. Metrics were computed by aggregating across participants and are scaling by max normalisation for ease of visualisation. These heatmaps demonstrate marked spatiotemporal variation in facial movement across expressions and conditions, providing direct evidence that the data are neither too homogeneous nor constrained by ceiling effects or simplistic dynamic patterns.

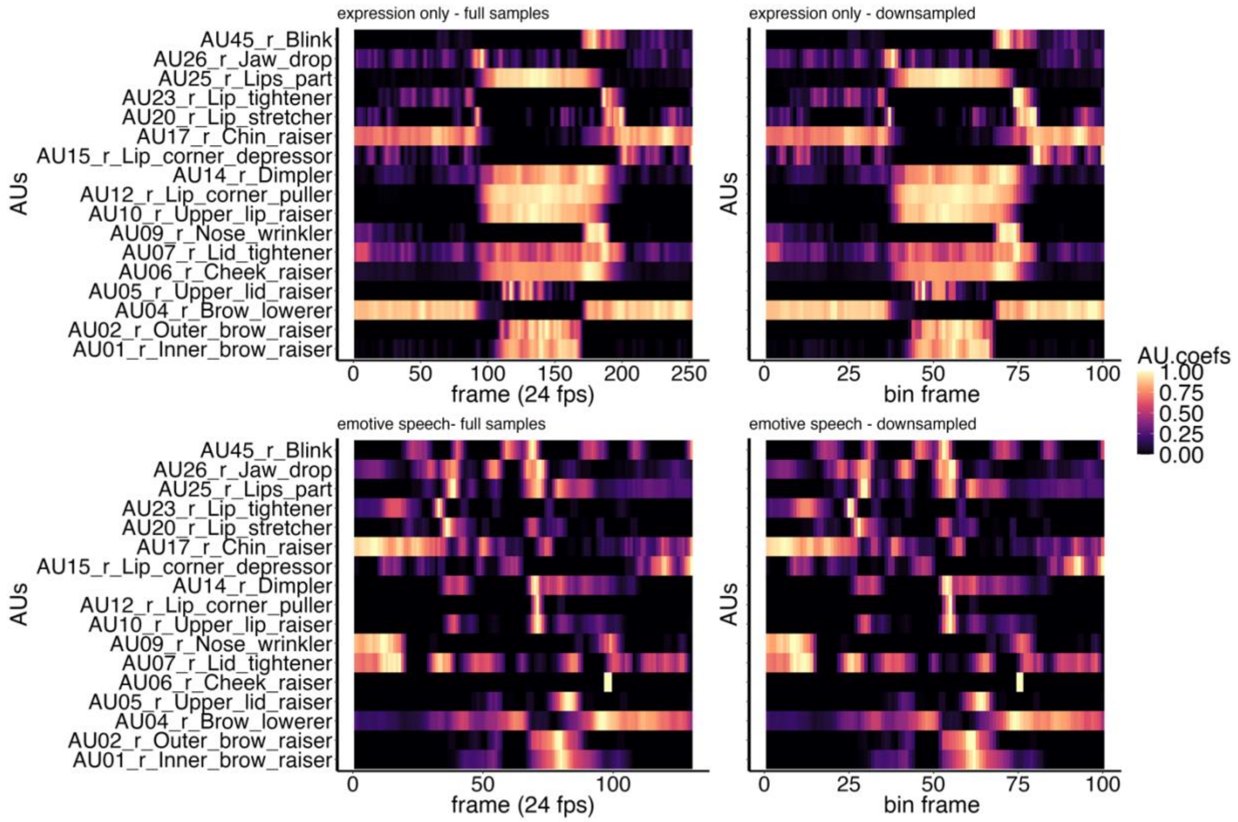


**Figure S11.** Comparison of full-sample (left) and down sampled (right) time-series for both *Expression Only* (top) and *Emotive Speech* (bottom) conditions. While video durations were largely consistent across stimuli, minor discrepancies were harmonised by down-sampling the facial Action Unit (AU) time-series into 100-time bins. We evaluated multiple thresholds and found that 100 bins provided a suitable balance, preserving the inherent variability needed to capture both the fast dynamics of emotive speech and the more stable patterns in expression-only segments.

**Table S5. List and brief description of timeseries features.**

| Feature name | Brief description |
|---|---|
| lempel_ziv | Measures the complexity of the series using Lempel-Ziv complexity. |
| approximation_entropy | Estimates the unpredictability (regularity) of the series. |
| sample_entropy | Computes the sample entropy, reflecting signal irregularity. |
| permutation_entropy | Uses the order relations of values to quantify complexity. |
| shannon_entropy_CS | Shannon entropy estimate (variant 1). |
| shannon_entropy_SG | Shannon entropy estimate (variant 2). |
| spectral_entropy | Entropy measure applied to the frequency domain. |
| nforbiden | Counts forbidden patterns, indicating complexity of sequences. |
| kurtosis | Measures the ‘tailedness’ of the distribution. |
| skewness | Gauges the asymmetry of the data distribution. |
| x_acf1 | First-order autocorrelation coefficient. |
| x_acf10 | Tenth-order autocorrelation coefficient. |
| diff1_acf1 | First-order autocorrelation of the first-differenced series. |
| diff1_acf10 | Tenth-order autocorrelation of the first-differenced series. |
| diff2_acf1 | First-order autocorrelation of the second-differenced series. |
| diff2_acf10 | Tenth-order autocorrelation of the second-differenced series. |
| x_pacf5 | Partial autocorrelation up to lag 5. |
| diff1x_pacf5 | Partial autocorrelation up to lag 5 of first-differenced data. |
| diff2x_pacf5 | Partial autocorrelation up to lag 5 of second-differenced data. |
| entropy | General entropy measure of complexity. |
| nonlinearity | Evaluates the degree of nonlinearity in the time-series. |
| hurst | Hurst exponent, indicating long-range dependence. |
| stability | Assesses changes in variance over time. |
| lumpiness | Captures fluctuations in variance over blocks of data. |
| unitroot_kpss | Tests stationarity via the KPSS test. |
| unitroot_pp | Tests stationarity using the Phillips-Perron test. |
| trend | Identifies a linear trend component. |
| spike | Detects sudden spikes or outliers. |
| linearity | Evaluates whether the underlying process is linear. |
| curvature | Identifies any curvature in the trend. |
| e_acf1 | Model residual autocorrelation at lag 1. |
| e_acf10 | Model residual autocorrelation at lag 10. |
| max_level_shift | Largest mean shift observed over the series. |
| time_level_shift | Time index where the maximum mean shift occurs. |
| max_var_shift | Largest variance shift in the series. |
| time_var_shift | Time index where the maximum variance shift occurs. |
| max_kl_shift | Largest Kullback-Leibler shift in distribution. |
| time_kl_shift | Time index of the maximum Kullback-Leibler shift. |

*Note*. The final analysis relied on 38 features only after dropping those that showed no variability or were unsuitable for our data (for instance, seasonality-based metrics) – for more details on the theoretical background and computation of these features see [3,4].
